# Molecular tracking devices quantify antigen distribution and archiving in the lymph node

**DOI:** 10.1101/2020.08.19.219527

**Authors:** Shannon M. Walsh, Ryan M. Sheridan, Thu Doan, Erin D. Lucas, Brian C. Ware, Rui Fu, Matthew A. Burchill, Jay R. Hesselberth, Beth A Jirón Tamburini

**Affiliations:** Department of Biochemistry and Molecular Genetics, University of Colorado School of Medicine, Aurora, CO, USA; Department of Medicine, Division of Gastroenterology and Hepatology, University of Colorado School of Medicine, Aurora, CO, USA; Immunology Graduate Program, University of Colorado School of Medicine, Aurora, CO, USA; RNA Bioscience Initiative, University of Colorado School of Medicine, Aurora, CO, USA; Department of Immunology and Microbiology, University of Colorado School of Medicine, Aurora, CO, USA

**Keywords:** Single-cell mRNA sequencing, lymph node, antigen processing, vaccination, antigen archiving, lymphatic endothelial cell, dendritic cell, nuclease resistant

## Abstract

Live, attenuated vaccines generate humoral and cellular immune memory, increasing the duration of protective immune memory. We previously found that antigens derived from vaccination or viral infection persist within lymphatic endothelial cells (LECs) beyond the clearance of the infection, a process we termed “antigen archiving”. Technical limitations of fluorescent labeling have precluded a complete picture of antigen archiving across cell types in the lymph node. We developed a “molecular tracking device” to follow the distribution, acquisition, and retention of antigen in the lymph node. We immunized mice with an antigen conjugated to a nuclease-resistant DNA tag and used single-cell mRNA sequencing to quantify its abundance in lymph node hematopoietic and non-hematopoietic cell types. At early and late time points after vaccination we found antigen acquisition by dendritic cell populations (DCs), associated expression of genes involved in DC activation and antigen processing, and antigen acquisition and archiving by LECs as well as unexpected stromal cell types. Variable antigen levels in LECs enabled the identification of caveolar endocytosis as a mechanism of antigen acquisition or retention. Molecular tracking devices enable new approaches to study dynamic tissue dissemination of antigens and identify new mechanisms of antigen acquisition and retention at cellular resolution *in vivo*.

## INTRODUCTION

Depending on the route of infection, vaccination mode, and ability of antigens to traffic, different dendritic cell (DC) subsets are required to initiate T cell priming. Upon subcutaneous immunization, small soluble proteins and virus particles pass through the lymphatics to the lymph node (LN), where LN-resident DCs acquire and present antigen^1, 2^. For larger antigens and/or pathogens that are too large to pass through the lymphatic capillaries, dermal DCs migrate to the LN for presentation of processed antigens to naïve T cells^1, 3, 4^. Most adaptive immune responses require antigen processing and presentation by conventional dendritic cells in either the draining lymph node or at the site of infection or vaccination (migratory cutaneous or dermal DCs) ^5^.

Previous studies have shown that viral antigens persist in the lymph node beyond the time frame of infectious virus ^6, 7, 8, 9, 10, 11^. We recently found that lymphatic endothelial cells (LEC) store antigens from viral infection and vaccination ^12, 13, 14^. Using a vaccine formulation that elicits robust cell-mediated immunity comprised of antigen, a Toll-like receptor (TLR) agonist, and an agonistic αCD40 antibody (TLR/αCD40 vaccination) or viral infection^15, 16, 17, 18, 19, 20, 21, 22, 23, 24, 25, 26^, we discovered that antigens were durably retained in the lymph node^12, 13, 14^. Antigen storage was dependent on the presence of a TLR agonist (e.g. polyI:C alone (TLR3/MDA5/RIGI or Pam3cys (TLR1/2)+ αCD40), but also occurred with antigen conjugated to a TLR agonist (e.g. 3M019 (TLR7)) ^14^. We named this process “antigen archiving” and showed it is important to poise memory T cells for future antigenic encounters^14^.

Prior to these studies, the only non-hematopoietic cell type thought to retain antigens were follicular dendritic cells, which harbor antigens in antigen-antibody complexes for extended periods of time and for the benefit of B cell memory ^11, 27^. Fibroblasts and non-endothelial stromal cells comprise a large portion of the lymph node stroma and are capable of presenting peripheral tissue antigens, but their capacity to acquire and present foreign antigens is not yet well understood ^28, 29, 30^. We were unable to detect antigen archiving by blood endothelial cells or fibroblasts in our initial studies^12, 13^. While LECs have been shown to present antigens in the absence of inflammation to induce T cell tolerance^31, 32, 33, 34, 35, 36, 37, 38^, we showed that presentation of archived antigen occurs only after exchange of the archived antigen from an LEC to a migratory DC; changing the stimulus from tolerizing to immunostimulatory^12, 13^. Soluble antigens are exchanged via two distinct mechanisms: (i) direct exchange between LECs and migratory DCs and (ii) LEC death. Antigen transfer from LECs to both migratory conventional (c)DC1s and cDC2s is required for archived antigen presentation to antigen-specific memory T cells ^12, 13^. After viral infection, archived antigen is transferred to *Batf3* dependent migratory DCs as a result of LEC death during lymph node contraction ^12^.

Limitations of current approaches have precluded sensitive and quantitative measures of antigen levels across cell types, providing only a glimpse of the cell types and molecular mechanisms that control antigen acquisition, processing and retention in the lymph node. Studies of antigen in the lymph node and peripheral tissues have mainly relied on antigen-flourophore conjugates or indirect measurement of antigen uptake and presentation^2, 6, 7, 11, 12, 14, 39^, which defined antigen acquisition by specific DC subsets and trafficking of antigens using live imaging^2^. However, antigen archiving has been difficult to study because antigen-fluorophore conjugates suffer from low microscopic detection sensitivity, yielding weak signals that diminish over time. Moreover, detection of antigen in the lymph node and other tissues has relied on flow cytometric analysis using cell surface markers, restricting analysis to specific cell types. To address these limitations and better understand antigen archiving, we developed a new approach to track antigen molecules throughout tissue-specific cell types *in vivo* using single-cell mRNA-sequencing.

## RESULTS

### Generation, validation, and immunogenecity of antigen-DNA conjugates

To quantify the dissemination and uptake of antigen in the draining lymph node (LN) after vaccination, we developed a strategy to measure antigen levels using single-cell mRNA sequencing. Many prior studies have used the model antigen, ovalbumin (ova), conjugated to a fluorophore to track antigen *in vivo*. Here we conjugated ova to DNA oligonucleotides with barcodes suitable for analysis by single-cell mRNA sequencing (**Fig. 1a**). The ~60 nt DNA tag contains a unique sequence barcode and PCR primer binding sites, similar to CITE-seq tags^40^ (**Supplementary Table 1**). We measured the stability of unconjugated DNA and ova-DNA conjugates in which the conjugated DNA either had normal phosphodiester linkages (pDNA) or was protected throughout by phosphorothioate linkages (psDNA). Quality control of these conjugates indicated a 1:1 stoichiometry of protein to DNA (**Fig. 1b**). To measure the stability of the anti-gen-DNA conjugate we added antigen-DNA conjugates to cultures of bone marrow derived dendritic cells (BMDCs) and quantified the amount of DNA in cell lysates and media over time. We found significantly higher levels of ova-psDNA in cells relative to ova-pDNA (~4-fold at day 1; *p* = 0.002 and ~7 fold at day 3; *p* = 0.004), indicating that psDNA is more stable than pDNA, but is processed by BMDCs at the same rate as ova protein (**Fig. 1c**). In addition, ova conjugation was required for phagocytosis by BMDCs, as we detected limited amounts of unconjugated pDNA or psDNA (values <1 at days 1-7) (**Fig. 1c**). We also measured conjugate stability in mouse LECs, a cell type that retains foreign proteins for long periods^14^, and found that ova-psDNA conjugates were stable over 7 days of culture, whereas ova-pDNA was rapidly degraded (**Fig. 1d**). As this was a measurement of the amount of DNA present we asked whether an ova protein-fluorophore conjugate was present over the same time frame as the ova-psDNA. Consistent with the detection of the psDNA, but not pDNA, we can detect ova protein by fluorescence in cultured LECs during the same time frame we were able to detect the psDNA (**Supplementary Fig. 1**).

**FIGURE 1.**
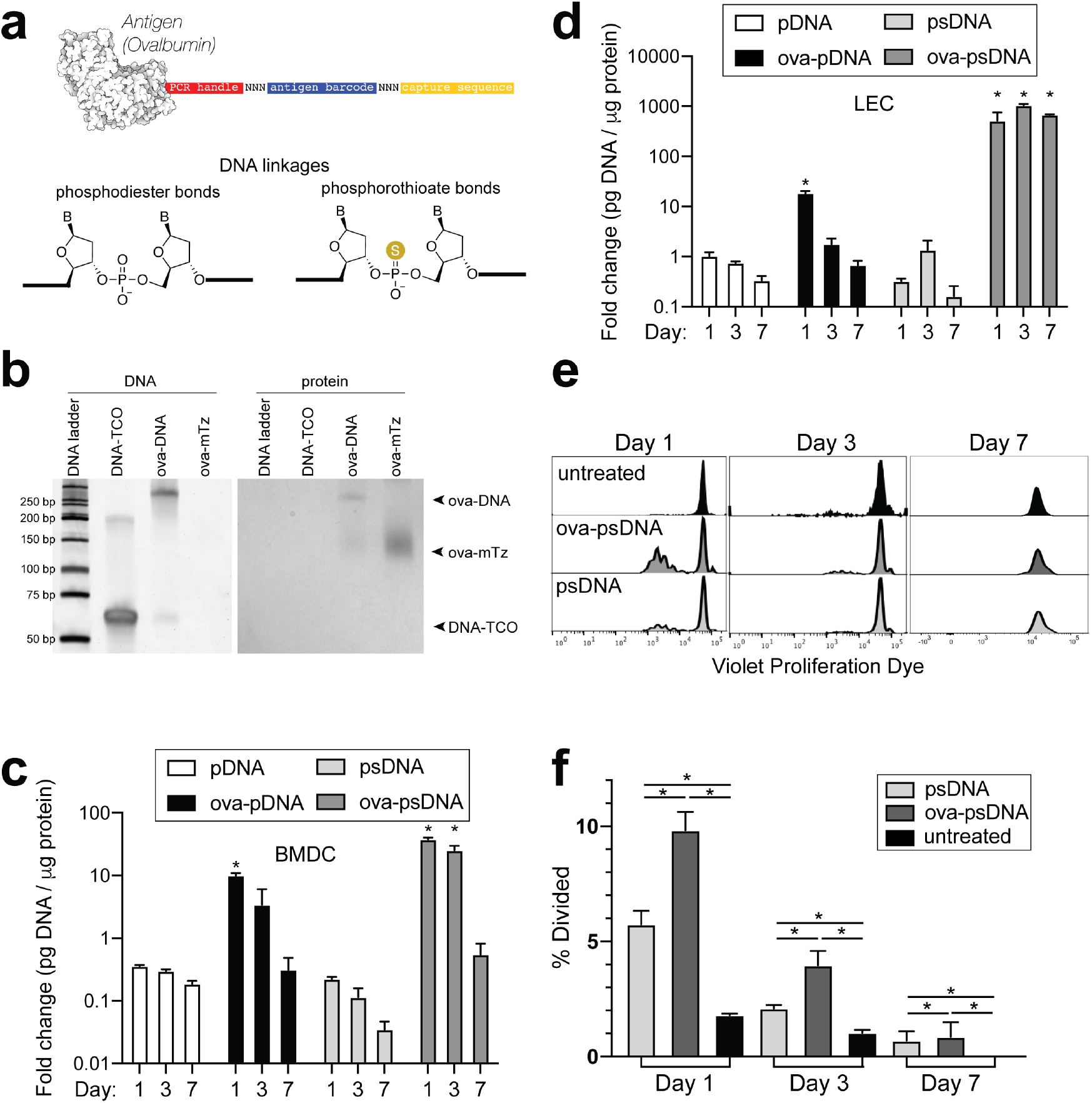
Antigen-psDNA conjugates undergo normal processing and presentation. **a**. Schematic of ovalbumin (PDB code 1ova) antigen conjugation to barcoded DNA with phosphodiester and phosphorothioate DNA linkages. Sulfur replaces a non-bridging oxygen to create a DNA phosphorothioate linkage. **b**. Conjugation of oligonucleotides to ovalbumin. Purified conjugate was analyzed by 10% TBE native PAGE stained with GelRed for DNA (left) followed by Coomassie staining for protein (right). DNA-TCO=61nt barcoded oligonucleotide with 5’-trans-cyclooctene (TCO); ova-mTZ=ovalbumin functionalized with methyltetrazine (mTZ); ova-DNA=DNA-conjugated ovalbumin product with oligonucleotide attached. **c**. Bone marrow derived dendritic cells (BMDC) were treated with pDNA, psDNA, ova-pDNA or ova-psDNA (5 μg) by addition to the culture media. After 1, 3, and 7 days cells were washed, released, lysed and analyzed for pDNA or psDNA by qPCR. Values are displayed as fold-change relative to the negative control (cells alone). Asterisks denote sample significant amounts relative to the negative control (*P* < 0.01; Wilcoxon rank-sum test). **d**. Analysis of DNAs as *in* (**c**) using murine lymph node lymphatic endothelial cells. **e**. BMDCs were incubated with ova-psDNA (conjugated), ova plus psDNA (unconjugated), or PBS for 1, 3, 7 days prior to adding OT-1 T cells labeled with violet proliferation dye. T cells and BMDCs were co-cultured at a ratio of 1:10 for 3 days. **f**. Quantification of **e** using the percent divided calculation described in the methods. Experiments were performed three times with 3-5 wells per sample with similar results. Asterisks denote sample significant amounts relative to the negative control (*P* < 0.05 Wilcoxon rank-sum test). Exact p-values are: Day 1 psDNA:ova-psDNA p=0.008, psDNA:untreated p=0.016, ova-psDNA:untreated p=0.016; Day 3 psDNA:ova-psDNA p=0.008, psDNA:untreated p=0.016, ova-psDNA:untreated p=0.016; Day 7 psDNA:ova-psDNA p=1, psDNA:untreated p=0.400, ova-psDNA:untreated p=0.400.

To determine whether conjugation of psDNA to ovalbumin affected ovalbumin processing and presentation, we measured BMDC presentation of ova-derived SIINFEKL peptide by co-culture with SIINFEKL-specific OT1 T cells. BMDCs given ova-psDNA induced significantly more proliferation of OT1 T cells than unconjugated ovalbumin (**Fig. 1e, f**) suggesting enhanced activation of BMDCs upon encounter with ova-psDNA conjugates. Furthermore, we detected pDNA and psDNA in BMDC culture media at one day after addition but not at later time points, confirming that ova-psDNA conjugates are processed and not released by BMDCs after phagocytosis (**Supplementary Fig. 2**). Finally, ova-psDNA conjugates led to increased OT1 proliferation at day 3 relative to ova plus psDNA (unconjugated), showing that ova-psDNA conjugates are immunostimulatory (**Fig. 1e,f**) and consistent with studies showing conjugation of antigens to RNA or DNA induce TLR7 (RNA) or TLR9 (DNA) signals that lead to prolonged antigen presentation ^41^.

We next asked whether vaccination with ova-psDNA conjugates elicits a T cell response *in vivo*. We compared antigen-specific T cell responses in mice vaccinated with a mixture of ova-psDNA and polyI:C/αCD40 to its individual components (ova, psDNA, polyI:C and polyI:C/αCD40; **Fig. 2a** and **Supplementary Fig. 3a**) and—consistent with the differences in OT1 proliferation we saw *in vitro*—found that T cell responses to ova-psDNA were greater than either ova with polyI:C, ova with polyI:C/αCD40, or a mixture of unconjugated ova and psDNA (**Fig. 2b**). Interestingly, ova-psDNA conjugate combined with polyI:C/αCD40 did not significantly enhance the T cell response beyond ova-psDNA alone (**Fig. 2b**). T cells stimulated by ova-psDNA produced significantly more IFNγ than any other vaccination strategy even in the absence of *ex vivo* SIINFEKL peptide stimulation, indicating prolonged and active presentation of ova-psDNA (**Fig. 2c,d**). Together these data show that ova-psDNA conjugates elicit antigenspecific T cell responses independent of polyI:C/αCD40. These findings are consistent with TLR9-dependent immune responses elicited by psDNA ^42, 43, 44^, similar to DC presentation of conjugates of ova demonstrated with other TLR agonists ^45^ and other subcutaneously administered ova-TLR conjugate vaccine platforms ^41^.

**FIGURE 2.**
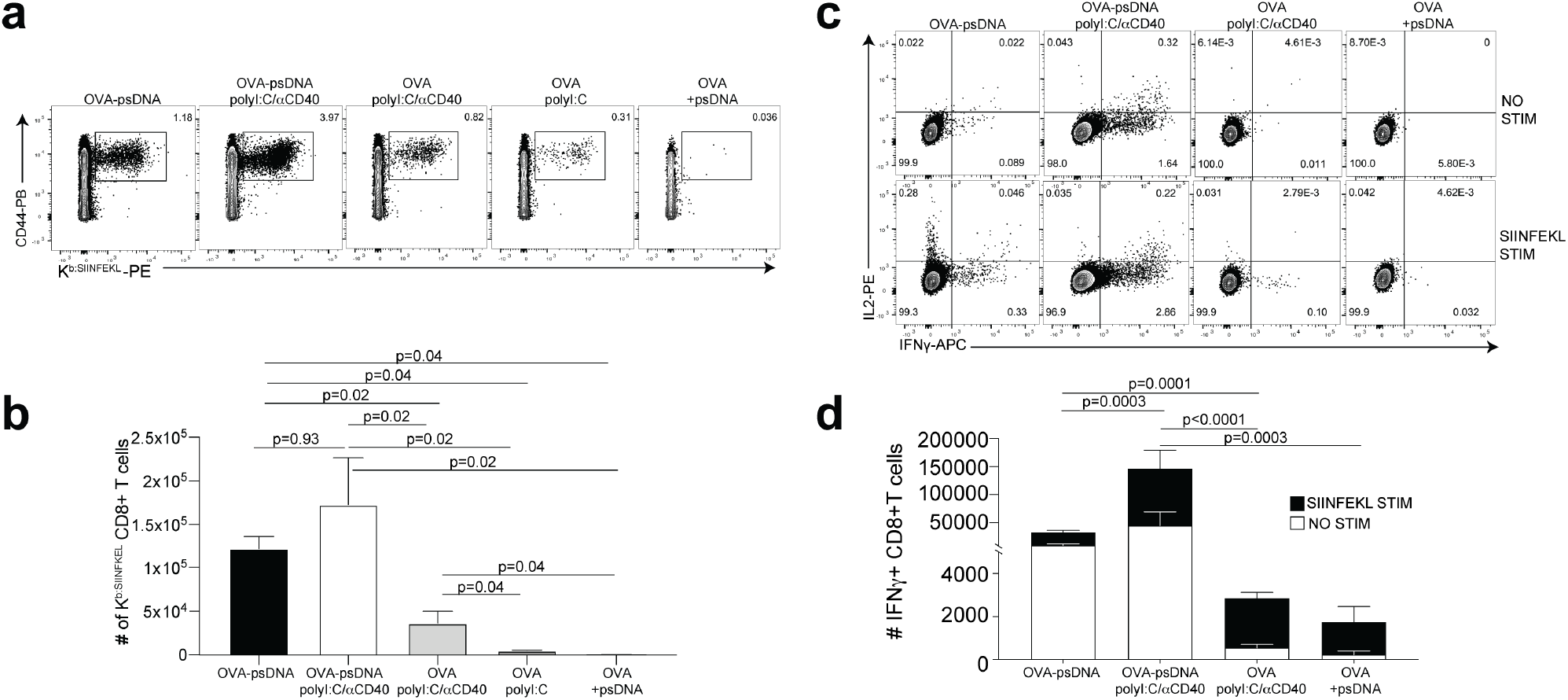
Antigen-psDNA conjugates elicit a robust immune response in vivo. **a**. Mice were immunized in the footpad with ova alone or ova-psDNA with or without polyI:C/aCD40 or polyI:C. After seven days, draining popliteal LNs were harvested and cells were stained and gated as B220-, CD3+, CD8+, CD44+ and OVA257 Kb SIINFEKL specific tetramer to measure antigen specific CD8 T cell responses. **b**. Quantification of SIINFEKL specific CD8 T cells within the lymph node (data from **a**). **c**. As in (**a**) and (**b**) except cells were restimulated with SIINFEKL peptide for 6 hrs ex vivo in the presence of brefeldin A, then stained for IFNγ and IL-2. **d**. Quantiation of interferon gamma positive CD8+ T cells with or without peptide stimulation in the draining lymph node. Experiment was performed 3 times; shown is combined data from at least 3 mice per group, per experiment. P-values were calculated using a two stage step up method of Benjamini, Krieger and Yekutieli and did not assume consistent standard deviation.

We previously showed that a vaccination strategy comprised of soluble antigen and vaccinia virus (Western Reserve; VV) induced robust antigen archiving that lasts longer than those using polyI:C/αCD40 adjuvant^12^. To evaluate antigen-psDNA performance during an active infection, we determined T cell responses after vaccination by comparing individual components with mixtures of ova, VV, ova-pDNA, or ova-psDNA. Subcutaneously administered ova-psDNA alone again elicited a T cell response (**Fig. 2** and **Supplementary Fig. 4a**) and addition of VV to ova-psDNA conjugate moderately increased T cell responses compared to ova-psDNA alone, similar to what we observed with ova-psDNA/polyI:C/αCD40 (**Fig. 2** and **Supplementary Fig. 4b**). Finally, we examined the cell type-specificity of ova-psDNA dissemination *in vivo*. Mice were vaccinated with mixtures of (i) ova-psDNA and VV or (ii) ova-psDNA and polyI:C/αCD40, and levels of ova-psDNA were quantified by PCR in both leukocytes and stromal cells (fractionated by CD45 expression) in the draining lymph nodes. We found that CD45-stromal cells had high amounts of ova-psDNA, corresponding to increased inflammation ^14^, whereas CD45+ leukocytes had very low levels of ova-psDNA 20 days after vaccination/infection (**Supplementary Fig. 4c**). These data recapitulate our previous demonstration of durable antigen retention by CD45-stromal cells^12, 14^, confirming that ova-psDNA is a faithful tracking device for antigen archiving *in vivo*.

### Molecular tracking of antigen during the immune reponse to vaccination

Given the ability of the antigen-psDNA conjugates to induce a robust immune response *in vivo* (**Fig. 2**) and our ability to use the psDNA as a measure of protein antigen levels (**Fig. 1**), we used the antigen-psDNA conjugates as a “molecular tracking device” to understand the distribution of the protein antigen in the lymph node following vaccination. To determine whether we could identify cells that acquire and archive antigens ^14^, we vaccinated mice subcutaneously with an equimolar mixture of uniquely barcoded ova-psDNA conjugate, unconjugated psDNA, and unconjugated pDNA (unprotected phosphodiester backbone) with VV (as in **Supplementary Fig. 4c**), and evaluated antigen distribution (via psDNA abundance) in the lymph node at early (2 days) and late (14 days) time points. At each time point, single-cell suspensions were prepared from draining popliteal lymph nodes and divided into stromal cell (by depleting CD45+ cells) or lymphocyte populations (by flow sorting for CD11c, CD11b, and B220 markers; **Supplementary Fig. 3b**). To enrich for myeloid cell populations but maintain representation of other cell types; CD11c+, CD11b+, B220+, and ungated live cells were mixed at a 4:4:1:1 ratio, respectively. These cell populations were analyzed by single-cell mRNA sequencing, measuring both mRNA expression and the quantity of psDNA in each cell using unique molecular identifiers ^46^ (**Fig. 3**).

**FIGURE 3.**
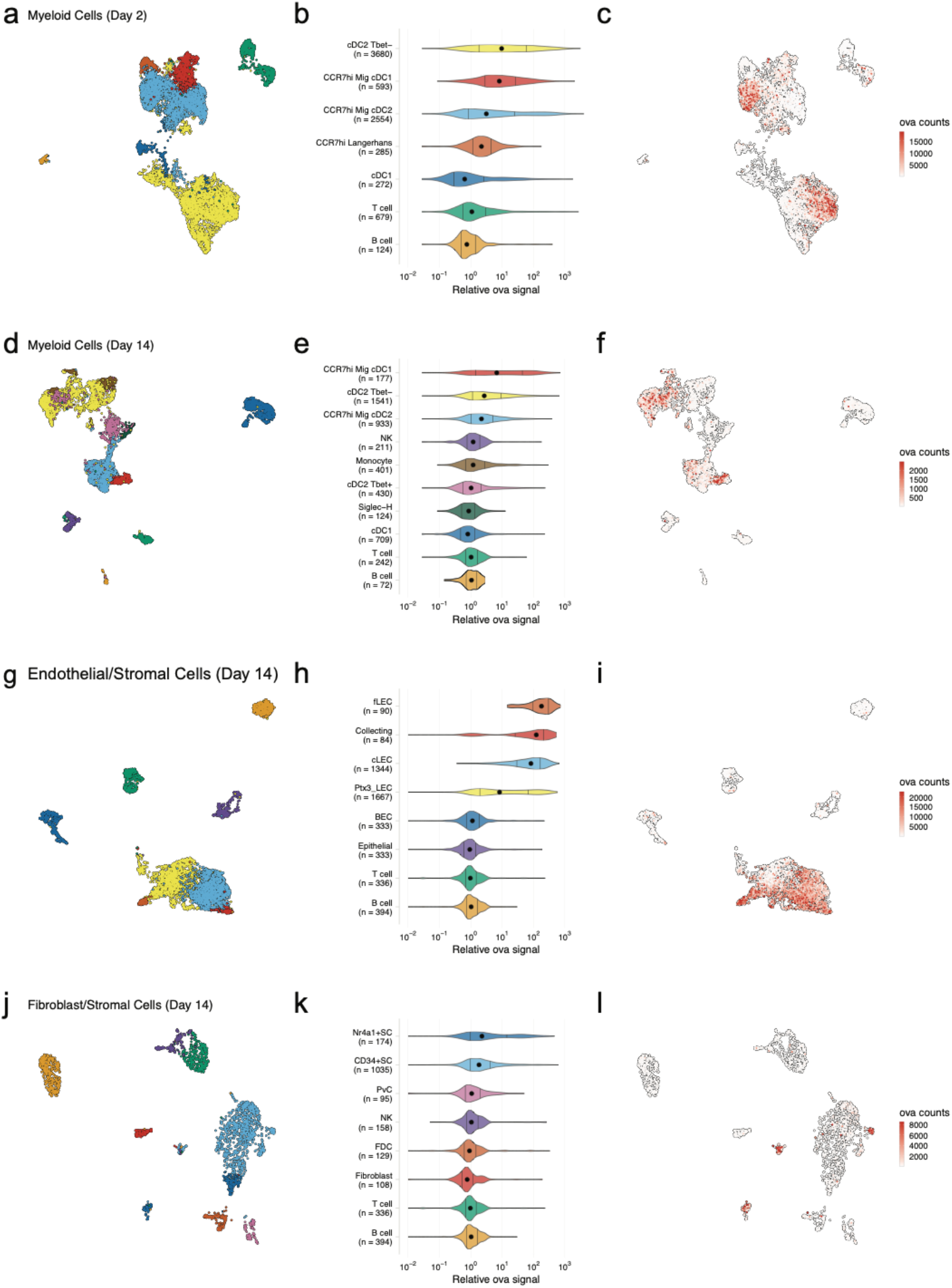
Dynamic acquisition of antigen-psDNA conjugates in lymph node tissue. (**a, d, g, j**) UMAP projections are shown for DCs (**a, d**), LECs (**g**), and FRCs (**j**) at day 2 (**a**) and day 14 (**d, g, j**). (**b, e, h, k**) Relative ova signal was calculated by dividing antigen counts for each cell by the median antigen counts for T and B cells. Signals are plotted on log_10_ scale; black dots indicate median values and vertical lines denote quartiles. Statistical comparisons between each pair of groups is available in **Supplemental Table 2**. (**c, f, i, l**) UMI-adjusted antigen counts are displayed on UMAP projections for each cell type.

We recovered a total of 800 cells in the CD45-fraction and 8,187 cells in the CD45+ fraction at the 2 day time point. We recovered more CD45-cells (6,372 CD45-; 4,840 CD45+) at the 14 day time point likely due to expansion and proliferation of the lymph node stroma^14, 22, 47^. We classified cell types using an automated approach^48^, comparing measured mRNA expression patterns to reference data sets for DCs^49, 50^, FRCs^51^, and LECs^52, 53, 54^ (**Supplementary Table 2**). As expected, the CD45+ fraction contained DCs, monocytes, T cells, and B cells (**Fig. 3a, b, d, e**), while the CD45-fraction contained stromal cells (SC), including LECs, blood endothelial cells (BEC), epithelial cells, and fibroblasts (**Fig. 3g, h, j, k**). We did not recover vaccinia viral mRNAs in cells at either time point, possibly due to viral clearance or a failure to recover infected, apoptotic cells in the live/dead selection (**Supplementary Fig. 3b**).

We first examined the dynamic changes of myeloid populations in the lymph node. We detected conventional DCs, including cDC1 and cDC2 (**Fig. 3a-c**), which develop from a common DC precursor upon expression of FMS-like tyrosine kinase 3 ligand (Flt3L)^55^. Lymph node (LN) resident and migratory cDCs can be distinguished by expression of cell-type-specific transcription factors including basic leucine zipper transcription factor (Batf3) and interferon regulatory factor (IRF8) (cDC1) ^56, 57, 58^ or IRF4 and Notch (cDC2)^59, 60^. These cDC types are also typically classified based on expression of CD11c, Zbtb46, and chemokine XC receptor 1 (cDC1 are XCR1+, cDC2 are XCR1-)^55, 61^. cDC2s are further categorized as either Tbet-dependent and anti-inflammatory (cDC2A) or RORγt-dependent and pro-inflammatory (cDC2B)^49^.

As expected, at day 2 we identified a large population of LN resident cDC2B (cDC2 Tbet-) cells harboring ova-psDNA^49^. However, we did not find any cDC2A (cDC2 Tbet+) cells, consistent with their role in anti-inflammatory processes^49^. The myeloid populations contained CCR7^hi^ cDCs (n = 3,432; 42% of total), which we classified as migratory DCs. This migratory DC population included Langerhans cells (n = 285; 3.5% of total), migratory cDC1s (n = 593; 7.2% of total), and migratory cDC2s (n = 2,554; 31% of total)^50^, migrating from the dermis (**Fig. 3b**). At day 14, we identified a population of LN-resident cDC2 Tbet+ cells (**Fig. 3e**) consistent with resolution of the immune response^49^. As cDC2 Tbet+ cells are thought to be anti-inflammatory, these data suggest that the immune response is being quelled (**Fig. 3e**). We also found a group of Siglec-H+ DCs, a cDC progenitor population ^49^ (**Fig. 3d,e**).

Using unique barcodes, we quantified the amount of ova-psDNA, psDNA, and pDNA across cell types. Levels of ova-psDNA molecules spanned four orders of magnitude, ranging up to 10^4^ unique molecules and depending on the cell types and time point (**Figure 3c,f,i,l**). In contrast to the large range of ova-psDNA across cell types, unconjugated psDNA and pDNA were largely undetectable, indicating that antigen conjugation is required for cell acquisition (**Supplementary Fig. 5**). Consistent with our previous studies^12^, we did not detect antigen-psDNA at appreciable levels in T cells or B cells (**Fig. 3**) and because these cell types were captured in both our CD45- and CD45+ samples, we used their median antigen levels to normalize antigen counts in other cell types across captures. We considered the trivial case wherein variation in antigen levels is explained by total mRNA abundance; these variables are uncorrelated in stromal cell types and weakly correlated in cDC subtypes, possibly reflecting activation status (**Supplementary Fig. 6**).

At the early day 2 time point, LN resident cDC2s contained high levels of antigen-psDNA, consistent with studies of soluble antigens^2^ (**Fig. 3b,c**). In addition, we found significantly higher levels of antigen in cDC2 Tbet-, migratory CCR7^hi^ cDC2s, and migratory CCR7^hi^ cDC1s (**Fig. 3b,c; Supplementary Table 2**), with an average of ~7-fold more antigen than T/B cells. At the later time point, migratory cDC1 cells contained the most antigen, consistent with previous studies^12^ (**Fig. 3e**). In addition, Tbet- and CCR7^hi^ migratory cDC2s contained moderate levels of antigen, up to 3-fold more than T/B cells, but had lower amounts of antigen relative to day 2 (**Fig. 3b,c,e,f; Supplementary Table 2**). At the late time point, we did not detect significant amounts of antigen in LN resident cDC1s, Tbet+ cDC2s, Siglec-H+ cells, or monocytes (**Fig. 3e**).

We next examined antigen levels in the LN stromal cell populations (**Fig. 3g-l**). Endothelial cells in the lymph node are classified by their association with blood or lymphatic vasculature; both are required for circulation and trafficking of immune cells to the lymph node. The blood vasculature circulates naïve lymphocytes to the LN and the lymphatic vasculature transports immune cells from the peripheral tissue including dermal DCs and memory T cells. We used an automated approach^48^ that uses correlation between reference and measured gene expression profiles to assign unknown cell types to subtypes defined by previous studies. While strong correlation reflects a good match between reference and query profiles, high correlation between multiple reference LEC subtypes^52, 53, 54^, and changes in expression induced by antigen acquisition made definitive cell type assignments challenging (**Supplementary Fig. 7b-d**). Notwithstanding these issues, we classified LEC subsets based on the highest correlation values to reference cell types (**Supplementary Fig 7d,e**)^54^ and identified three LEC subtypes^52, 53, 54^ including Ptx3 LECs, ceiling LECs, and Marco LECs with high levels of antigen at the early time point (**Supplementary Fig. 7a**). At the late time point, expansion and proliferation of lymph node stromal cells contributed to larger populations of cells including floor LECs, collecting LECs, ceiling LECs, Ptx3 LECs^53^, and blood endothelial cells (BEC) (**Fig. 3h**)^62^.

At the day 14 time point, several LEC subtypes maintained high antigen levels (**Fig. 3h, Supplementary Table 2**). Floor LECs had uniformly high amounts of antigen, confirming our previous studies using flow cytometric analysis of fluorescent antigen^12^. Median levels of antigen in collecting, Ptx3, and ceiling LEC populations were significantly higher than B/T cells, but cells in these groups contained a range of antigen with both high and low populations. We hypothesized this variability stems from the physical location of the LECs within the LN and their access to trafficking antigen, and confirmed that fluorescent antigen amounts are highest on subcapsular LECs as identified by surface expression of PD-L1 and ICAM1 two weeks after immunization ^22, 32^ (**Supplementary Fig. 8**). Together our findings suggest that antigen first passes through the sinus followed by the cortex and medulla. These data also suggest that populations of LECs with less antigen could be a result of how the antigen travels through the lymph node or mechanisms of antigen release over time.

Similar to the endothelial cell population, the number and types of non-endothelial stromal cells increased at the later time point after immunization. Non-endothelial stromal cells in the lymph node are classified by their location in the lymph node into T-zone reticular cells (TRC), marginal reticular cells (MRC), follicular dendritic cells (FDC), and perivascular cells (PvC)^51^. Recently, additional subsets were identified including: Ccl19^lo^ TRCs located at the T-zone perimeter, Cxcl9^+^ TRCs found in both the T-zone and interfollicular region, CD34^+^ stromal cells found in the capsule and medullary vessel adventitia, indolethylamine N-methyltransferase^+^ stromal cells found in the medullary chords, and Nr4a1^+^ stromal cells^51^.

At the early time point, the Cxcl9+ TRCs and CD34+ SCs^51^, had high amounts of antigen (~10-fold relative to T/B cells) (**Supplementary Fig. 9** and **Supplementary Table 2**). At the late time point, we detected CD34+ SCs, Nr4a1+ SCs, follicular dendritic cells (FDC), and PvCs (**Fig. 3k**). Only the CD34+ and Nr4a1+ SCs contained significant amounts of antigen (**Fig. 3k, Supplementary Table 2**). Interestingly, the CD34+ stromal cells are adjacent to ceiling LECs and the Nr4a1+ SCs are found in the medullary chord and medullary sinus, which are lined by medullary LECs. These findings may suggest potential antigen exchange mechanisms between LECs and SCs that have yet to be defined. We found little antigen in PvCs or FDCs (**Fig. 3k**).

Finally, these data provided insight into antigen transfer between stromal and dendritic cells, a process important for enhanced protective immunity ^12, 14^. We previously showed that archived antigen is transferred from LECs to migratory *Batf3*-dependent cDC1s two weeks after infection^12^. Here we confirm that CCR7^hi^ migratory cDC1s had the highest amount of antigen two weeks after vaccinia infection (**Fig. 3e**)^12^. Together, these data validate the use of molecular tracking devices by corroborating previous studies of antigen trafficking and identify new cells types that dynamically acquire antigen during infection.

### Gene expression signatures associated with antigen acquisition by dendritic cells

We next leveraged the variation in antigen levels across cell types (**Fig. 3b,e,h,k**) to identify gene expression signatures associated with high levels of antigen that would validate our approach. We classified cells as “antigen-high” and “antigen-low” using a two-component mixture model, and identified marker genes associated with each class (**Fig. 4a,b**). To validate this approach, we evaluated the dendritic cell populations, as genes associated with phagocytosis and activation have been established^50, 63, 64, 65, 66, 67, 68, 69, 70, 71, 72^. Dendritic cell populations generally contained lower antigen levels that were variable across subtype (**Fig 3**). We classified antigen-low and antigen-high cells for each subtype. We found the highest antigen levels and largest differences in gene expression for Tbet-cDC2 cells (277 genes in antigen-high cells, **Fig 4, Supplementary Table 4**), consistent with cDC2s acting as the primary cell type of antigen uptake following protein immunization^2^.

**FIGURE 4.**
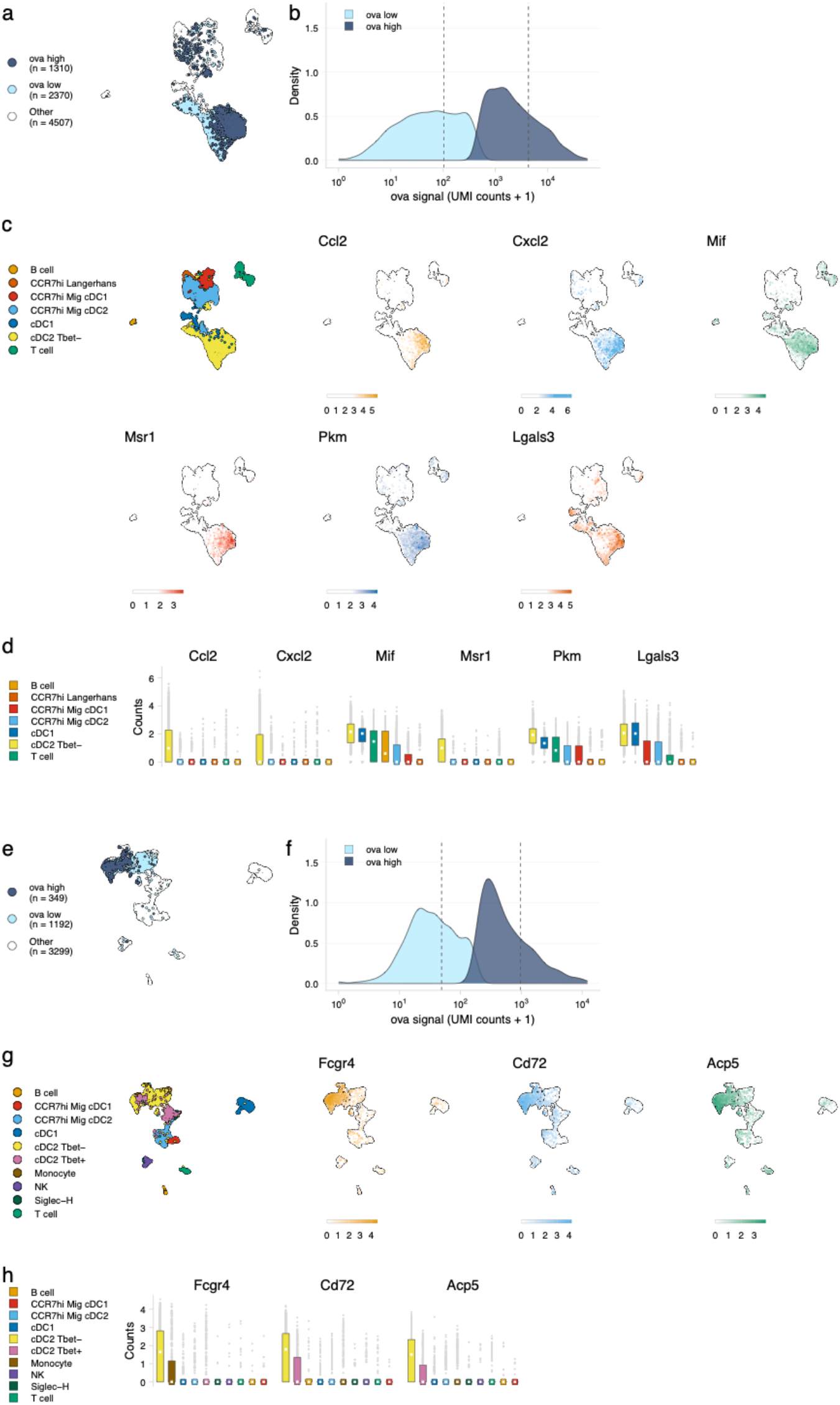
Antigen-based classification of DCs and validation of genes associated with DC activation. (**a, e**) Day 2 (**a**) and day 14 (e) cDC2 Tbet-cells containing low and high antigen counts were identified using a two component mixture model. A UMAP projection is shown for ova-low and ova-high cells. Cell types not included in the comparison are shown in white (other). (**b, f**) The distribution of ova antigen counts is shown for ova-low and ova-high cDC2 Tbet-cells. Dotted lines indicate the mean counts for each population. (**c, g**) UMAP projections show the expression (log-normalized counts) of top markers associated with ova-high cDC2 Tbet-cells. (**d, h**) Expression (log-normalized counts) of antigen-high markers in each cell type.

At the early time point, genes upregulated in antigen-high DCs confirmed DC activation (**Supplementary Table 4**). Antigen-high cDC2 Tbet-cells upregulated genes *Ccl2* and *Cxcl2* (consistent with active recruitment of inflammatory cells^66, 69^), *Msr1* (consistent with antigen scavenging^71^), as well as *Pkm, Galectin 3*, and *Mif* (consistent with DC-T cell responses and DC differentiation during inflammation^63, 65, 68^) (**Fig. 4c,d**).

At the late day 14 time point, the highest antigen counts were found in the migratory cDC1 population, consistent with a role for migratory cDC1s in archived antigen acquisition from LECs^12^ (**Fig. 3e,f**). Among the genes highly expressed by the antigen-high CCR7^hi^ migratory cDC1 population were *Ccl5* and *Fscn1* (**Supplementary Table 4**). Consistent with these DCs being involved in archived antigen presentation, *Ccl5/RANTES* regulates CD8 T cell responses during chronic viral infection^73^ and *Fscn1*, an actin binding protein, regulates cell migration of mature DCs via podosome formation^74^. Similar to the day 2 timepoint, the differences in gene expression for antigen high and low cells were greatest within for Tbet-cDC2 populations (230 genes in antigen-high cells; **Fig. 4e,f** and **Supplementary Table 4**). Genes upregulated in antigen-high tbet-cDC2s included *Fcgr4*, which is involved in phagocytosis, antigen presentation, and proinflammatory cytokine production^67, 70^, and *CD72* and *Acp5*, which are important for the inflammatory response and pathogen clearance^64, 72^ (**Fig. 4g,h**). Collectively, these genes evoke specific processes in DC subsets required for the immune response; it remains to be determined whether they are specifically associated with LEC-DC antigen exchange or storage of antigens within DCs.

### Gene expression signatures associated with antigen archival by LECs

We next evaluated the LEC population to determine whether our classification approach could identify genes involved in antigen archiving. We applied the classifier to LECs as a population and found large numbers of antigen-high-floor, collecting, and ceiling LECs (**Fig. 5c**). Ptx3 LECs were comprised of a mixture of antigen-low and antigen-high cells, but there was a larger fraction of Ptx3 LECs with low antigen (**Fig. 5c**). There were less antigen-low LECs compared to antigen-high LECs overall (34% of total), suggesting that antigen archiving may be specific to LECs in general rather than attributable to a specific LEC subset (**Fig. 5**).

**FIGURE 5.**
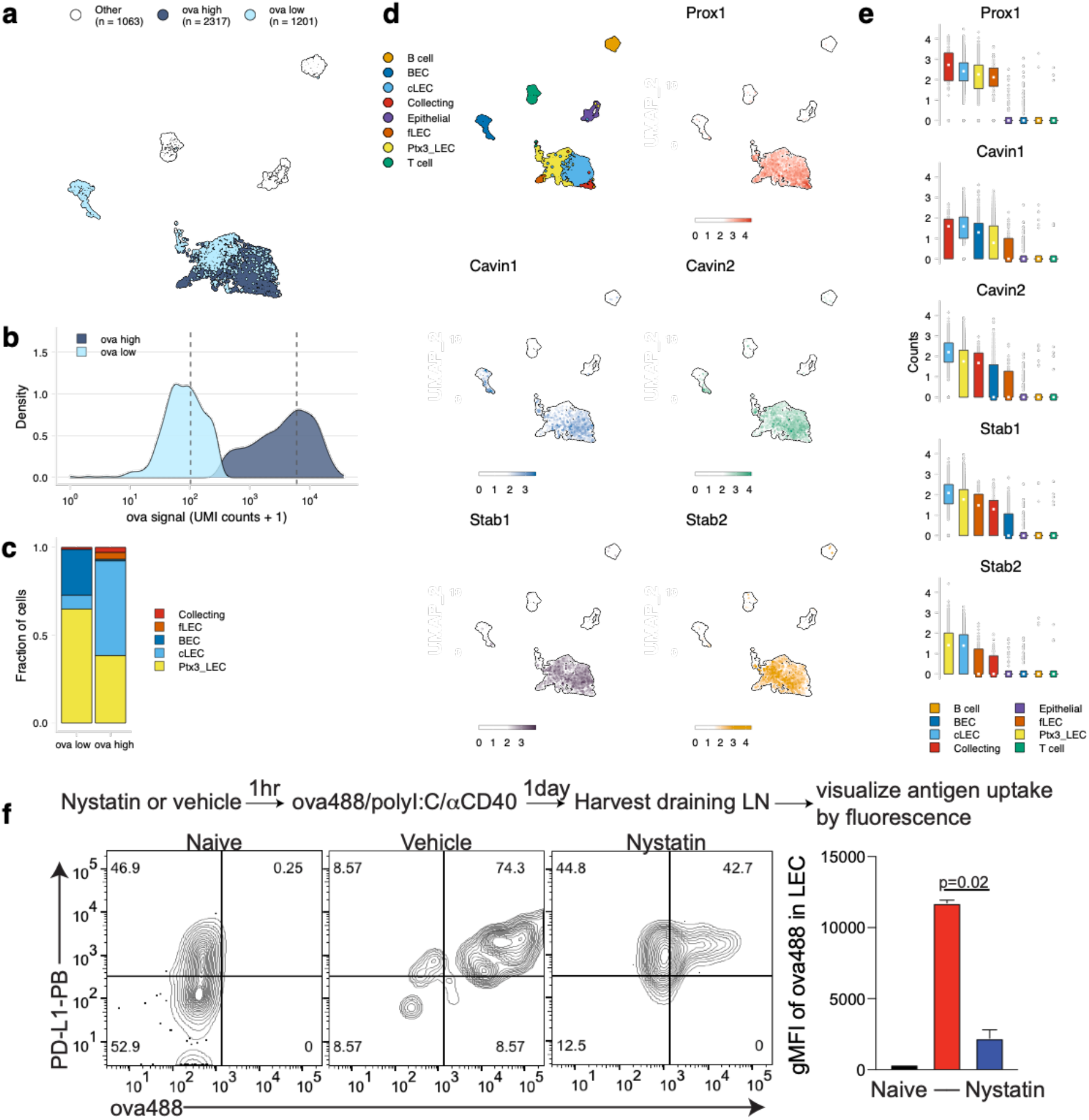
Antigen-based classification of LECs and identification of marker genes. **a**. Day 14 LECs were classified into antigen-high and antigen-low using a two-component Gaussian mixture model. A UMAP projection is shown for antigen-low and antigen-high cells. T cells, B cells, and epithelial cells are shown in white (Other). **b**. Distribution of antigen counts for antigen-low (light blue) and antigen-high (dark blue) cells. Dotted lines indicate mean counts for each population. **c**. The fraction of cells belonging to each LEC cell type for antigen-low and antigen-high populations. **d**. UMAP projections show expression of genes significantly enriched in the antigen-high population (scale is log-normalized counts). **e**. Expression (log-normalized counts) of antigen-high markers in each cell type. **f**. Mice were injected in the footpad with nystatin (dose) and 1 hour later ova488/polyI:C/αCD40. After 24 hours, mice were euthanized and draining popliteal LN removed, stained for LEC markers (CD45, PDPN, CD31, PDL1) and gated as in **Supplementary Fig. 8**. Shown are representative flow flots and quantification of geometric mean fluorescence intensity (gMFI) from naïve (black bar), vehicle control (red bar) and nystatin treated (blue bar). Three mice per group were evaluated and experiment was performed 3 independent times with similar results. Nystatin treatment reduces ova488 signal in LECs relative to vehicle (*P*=0.02; Wilcoxon rank-sum test).

Using this classification approach, we identified 142 mRNAs that were significantly changed in antigen-high or antigen-low LECs (**Supplementary table 3**). *Prox1*, while expressed by all LECs identified, was highly expressed in antigen-high LECs, independent of the LEC type (**Fig 5d, e**). *Prox1* is a transcription factor required for LEC differentiation from blood endothelial cells and defines LEC identity via regulation of *Vegfr3, Pdpn*, and *Lyve-1*^75, 76, 77^. *Prox1* upregulation in antigen-high LECs indicates it may also transcriptionally regulate processes involved in antigen archiving.

Upregulation of *Cavin1* and *Cavin2* by antigen-high LECs suggested that caveolar endocytosis may contribute to antigen acquisition by LECs, consistent with LEC dynamin-mediated transcytosis *in vitro* ^78^ (**Fig. 5d,e**). *Cavin2* appears more specific to LECs than *Cavin1*, which is also upregulated by BECs, suggesting that *Cavin2* mediates endocytosis specifically in endothelial cells of the lymphatic lineage. Based on *Cavin2* gene expression it appears this process may be most active in ceiling LECs (**Fig. 5e**). To confirm this finding, we asked whether inhibition of the caveolin pathway with nystatin impaired endocytosis of fluorescent antigen in mice vaccinated with ovalbumin/polyI:C/αCD40. We found a significant decrease in antigen acquired by LECs in the nystatin treatment group 24 hours after administration of fluorescent antigen with an innate immune stimulus (**Fig. 5f**), affirming the utility of molecular tracking devices for identifying genes involved in the process of antigen acquisition or archival.

Finally, expression of Stabilin-1 (*Stab1*) and Stabilin-2 (*Stab2*) is increased in antigen-high LN endothelial cells, suggesting that scavenging pathways are required for the acquisition of antigen-psDNA conjugates after vaccination. *Stab2* is uniquely expressed by LECs in the lymph node and not by BECs ^79^, and Stabilin-1 and Stabilin-2 act as receptors for internalization of antisense oligonucleotides with phosphorothioate linkages in liver endothelial cells and Kupffer cells ^80^. However, we did not find significant amounts of unconjugated psDNA in LECs (Supplementary Fig. 5), indicating that *Stab1/Stab2* are upregulated as part of an antigen scavenging or trafficking program initiated in LECs upon antigen acquisition during infection.

## DISCUSSION

Our development of a “molecular tracking device” enabled tracking of antigen throughout the lymph node to specific cell types that acquire and archive antigens following subcutaneous immunization. Previous studies used canonical surface markers to track antigen by microscopy and flow cytometry; instead, our approach simultanesously defines cell type by gene expression and quantifies the acquired antigen. The molecular tracking device will enable the study of archived antigens at time points beyond the lifetime of antigen-fluorophore conjugates and provided of a more complete catalog of cell types involved in antigen acquisition and retention.

Our approach validates and expands upon our previous studies of antigen archiving and cell types that enhance protective immunity. Both here and in our previous studies we found that whereas LECs archive antigen, migratory DCs passing through the lymphatic vasculature are required to retrieve and present archived antigen to memory CD8 T cells derived from the initial infection or immunization^5^. Antigen exchange from LECs to DCs and subsequent DC presentation yields memory CD8 T cells with robust effector function during infectious challenge. Several recent reports defined LEC and non-endothelial stromal cell subsets within the lymph node^51, 52, 53, 54^. By combining our molecular tracking device with these reference cell types, we found that non-endothelial stromal cell types acquire foreign antigens including CD34+ stromal cells, which neighbor subcapsular sinus LECs in the tissue ^51^. These findings suggest that the interstitial pressure created by subcutaneous vaccination allows antigens to pass through the tissue directly to the LN capsule, bypassing the lymphatic capillaries. Intriguingly, bypass of lymphatic capillaries may still lead to LEC acquisition of antigens from the CD34+ stromal cells via SC-LEC exchange. Such a mechanism would encourage future LEC-DC interactions and provide a benefit to protective immunity.

Molecular tracking devices provide a measure of cell state orthogonal to gene expression, which we leveraged to identify candidate pathways involved in antigen acquisition (**Fig. 4**). We show that the caveolin pathway is upregulated in antigen-high LECs and confirmed this pathway is involved in antigen acquisition *in vivo* following vaccination via pharmacological inhibition of caveolar endocytosis (**Fig. 4f**). Genes uniquely expressed by LECs such as *Prox1, Cavin2* and *Stab2*^50, 79, 81^ represent targets for further manipulation of antigen archiving by LECs.

The psDNA component of the tracking device elicits an immune response similar to other TLR-antigen conjugate vaccines^82, 83^, likely due antigen-psDNA stability within dendritic cells that causes prolonged antigen presentation in the cells that acquire the antigen^45, 41^. This effect is illustrated by increased IFNγ production in the absence of *ex vivo* peptide stimulus (ova-psDNA compared to unconjugated ova; **Fig. 2**). Prolonged antigen presentation better replicates an infection wherein levels of viral or bacterial antigen rise over the duration of infection. However, in other applications it may be helpful to limit the immunoreactivity of the antigen-psDNA via cytosine methylation^84^ or backbone modification^85^.

Molecular tracking devices will enable new approaches to study molecular dissemination *in vivo*. To date, protein-DNA conjugates have been deployed in single-cell mRNA sequencing experiments for *ex vivo* staining applications (e.g., CITE-seq ^46^). Our study lays the groundwork for molecular tracking devices involving protein, antibody, drug, or pathogens conjugated to nuclease-resistant, barcoded oligonucleotides that are stable during transit through mouse tissues. The approach naturally extends to understanding how multiple different antigens might be processed (using unique DNA barcodes) and enables new studies to manipulate antigen archiving to improve vaccines, vaccine formulations, and prime-boost strategies. Moreover, the oligonucleotide portion of the tracking device should enable analysis of its distribution in cells by in situ hybridization or intact tissue by spatial transcriptomics^86, 87, 88^, obviating the need for antibody-mediated detection of antigen.

## MATERIALS AND METHODS

### Mice

4-6 week old mice were purchased from Charles River or Jackson Laboratory, unless otherwise stated, bred and housed in the University of Colorado Anschutz Medical Campus Animal Barrier Facility. Wild type and OT1 mice were all bred on a C57BL/6 background. OT1 mice are a TCR transgenic strain specific to the SIINFEKL peptide of ovalbumin (OVA257-264) in the context of H-2K^b^. All animal procedures were approved by the Institutional Animal Care and Use Committee at the University of Colorado.

### Phosphorothioate and phosphodiester oligonucleotides

Oligonucleotides were synthesized by Integrated DNA Technologies (IDT) and contained a 5’ amine for conjugation, primer binding site, barcode, 10xGenomics Gel Bead Primer binding site for capture sequence 2, and a 3’ biotin. Phosphorothioated oligonucleotides contained a phosphorothioate modification at every linkage. All oligonucleotide sequences can be found in **Supplementary Table 1**.

### Conjugation of oligonucleotides to protein

Oligonucleotides were conjugated to ovalbumin by iEDDA-click chemistry^89^. Oligonucleotides were derivatized with trans-cyclooctene (TCO) in 10X borate buffered saline (BBS; 0.5 M borate, 1.5 M NaCl, pH 7.6; sterile filtered). Dilution of this buffer to 1X results in a final pH of 8.5. A reaction containing 40 nmol of amine-modified oligo (0.5 mM), 1X BBS, 10% DMSO, 8 μL of 100 mM TCO-PEG4-NHS in DMSO (10 mM final; Click Chemistry Tools, A137), pH 8.5 was rotated at room temperature for 15 min. A second aliquot containing the same amount of TCO-PEG4-NHS in DMSO was added and the reaction was rotated at room temperature for another hour. Excess NHS was quenched by adding glycine, pH 8.5 to a final concentration of 20 mM and rotated at room temperature for 5 min. Modification was confirmed by analysis on an 8% denaturing TBE PAGE gel. Samples were precipitated by splitting the reaction into 20 μL aliquots and adding 280 μL of nuclease-free water, 30 μL of 3 M NaCl, and 990 μL of 100% ethanol. The precipitation reaction was incubated at −80 °C overnight followed by centrifugation at >10,000 *x g* for 30 min. The supernatant was discarded, the pellet was washed with 100 μL of 75% ethanol, and centrifuged at >10,000 *xg* for 10 min. The supernatant was removed, and the pellets were dried for 5 min at room temperature. The pellets were recombined by resuspension in 50 μL of 1X BBS. Samples were quantified by *A*_260_.

To conjugate methyltetrazine to ovalbumin: detoxified Ovalbumin (Sigma-Aldrich, St. Louis, MO) (using a Triton X-114 lipopolysaccharide detoxification method^90^), was buffer exchanged into 1X BBS, pH 8.5. To an Amicon 0.5 mL 30 kDa filter (Millipore, UFC5030) was added 1 mg of ovalbumin and 1X BBS to a volume of 450 uL. The filter was centrifuged at 14,000 *xg* for 5 min. The flow through was discarded and the sample washed twice with 400 μL of 1X BBS. The product-containing column was inverted into a clean collection tube and centrifuged at 1,000 *xg* for 2 min. Assuming no loss, the volume of the sample was adjusted to 2 mg/mL with 1X BBS. 400 μL of 1X BBS was added to the Amicon filter and stored at 4 °C for later use. A 500 μL labeling reaction containing 0.5 mg of ovalbumin in 1X BBS and 50 μL of 2 mM mTz-PEG4-NHS in DMSO (0.2 mM final; Click Chemistry Tools, 1069), pH 8.5 was rotated at 4 °C overnight. Excess NHS was quenched by adding glycine, pH 8.5 to a final concentration of 20 mM and rotated at room temperature for 10 min. The previously stored Amicon filter was centrifuged at 14,000 *xg* for 5 min and the flow through discarded. 400 μL of reaction mixture was added to the filter and centrifuged at 14,000 *xg* for 5 min. This was repeated until all 1 mg of protein had been added to the filter and was supplemented with 1X BBS as needed. Samples were washed 1X with 400 μL of 1X BBS. The product-containing column was inverted into a clean collection tube and centrifuged at 1,000 *xg* for 2 min. Assuming no loss, the volume of the sample was adjusted to 5 mg/mL with 1X BBS.

For the final antigen-DNA conjugation, a 100 μL reaction containing 300 μg of ovalbu-min-mTz and 6 nmol of oligonucleotide-TCO (1:1 equivalents) in 1X BBS was rotated at 4 °C overnight. Excess mTz was quenched with 10 μL of 10 mM TCO-PEG4-glycine and rotated at room temperature for 10 min. TCO-PEG4-glycine was prepared by reaction of 10 mM TCO-PEG4-NHS with 20 mM glycine, pH 8.5 in 1X BBS for 1 h at room temperature and stored at −20 °C. Products were analyzed by 10% TBE PAGE. For purification, *e*xcess ovalbumin and DNA were removed by filter centrifugation. 200 μL of 1X PBS was added to an Amicon 0.5 mL 50 kDa filter (Millipore, UFC5050) followed by 300 μL of sample. The filter was centrifuged at 14,000 *xg* for 5 min and the flow through discarded. Samples were washed five times with 400 μL of 1X PBS and centrifuged at 14,000 *xg* for 5 min. The product-containing column was inverted into a clean collection tube and centrifuged at 1,000 *xg* for 2 min. Purified products were analyzed by 10% TBE PAGE and total protein quantified with Bio-Rad protein quantification reagent (Bio-Rad, 5000006). LPS contamination after conjugation was below 0.5 EU/mg as mentioned below in the vaccinations section.

### Bone marrow derived dendritic cell and lymphatic endothelial cell cultures

Both left and right tibia and femur were isolated under sterile conditions. Bone marrow was extracted from femurs of 6-8-week-old C57BL/6 mice by decollating the top and bottom of the bone and releasing the marrow with 27 gauge syringe and 5ml of Modified Essential Medium (Cellgro). Suspension was strained through 100um filter, pressed with the back of a syringe and washed. Cells were spun 1500RPM, 5 min then suspended in MEM with 10% FBS, 20ng/ml of GM-CSF from the supernatant of the B78hi-GM-CSF cell line. Every 2 days, dead cellular debris was spun, supernatant collected and combined 1:1 with new 40ng/ml GM-CSF 20% FBS (2x) in MEM. After 7 days of culturing at 37C, 5% CO2, cells were harvested for respective assays. Mouse lymphatic endothelial cells (Cell Biologics, C57-6092) were cultured in Endothelial Cell Media (Cell Biologics, M1168) with kit supplement. T75 Flasks were coated with gelatin (needs Cat#) for 30 minutes 37C, washed with PBS and then inoculated with mLEC. Cells were passaged with passive trypsin (Cat#) no more than 3-6 times and split at density of 1:2.

### Conjugate detection assay

BMDC and mLEC Cultures were stimulated with 20 μg of anti-CD40, 20 μg Poly I:C, and 5 μg of either OVA-psDNA or OVA in a 6-well format. 24hr post treatment, cells were washed and refreshed with new media. At designated time points, cells were harvested, counted and transferred into micro-centrifuge tubes, spun at 350g, and both supernatant and pellets were frozen at −80C. Cell pellets were lysed in 50 μL of Mammalian Protein Extraction Reagent (MPER; Thermo Scientific, 78503). Conjugate DNA was measured by qPCR amplification from 1 μL of lysate in a 10 μL reaction containing 5 μL of iTaq Universal SYBR Green Supermix (Bio-Rad, 1725125) and 5 pmol of each primer (**Supplementary Table 1**). Quantification was measured using an external standard curve and normalized to lysate protein content.

### OT1 Isolation and co-culture

CD8 T cells were isolated from an OT1+ mouse using the mojosort mouse CD8 T cell isolation kit (Biolegend) and labeled with violet proliferation dye (BD Biosciences cat# 562158). For DC-T cell co-culture, BMDCs were treated with psOVA (5μg), or OVA+psDNA (5μg) for 1,3 or 7 days. BMDCs were washed and then co-cultured with labeled OT1s for three days at a 1:10 ratio of BMDC:OT1. Cells were then stained and run on a flow cytometer. OT1 division (percent dividing cells) was calculated as previously described^91^ using the equation fraction 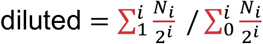 where *i* is the generation number (0 is the undivided population), and N*_i_* is the number of events in generation *i*.

### Vaccinations

6-8 week-old C57BL/6 (CD45.2) mice were immunized with 1E3 or 1E4 colony forming units of Vaccinia Western Reserve or 5 μg of Poly I:C (Invivogen) with or without 5μgof anti-CD40 (FGK4.5, BioXcell)and 10 μg åof OVA-psDNA or OVA in 50ul volume by footpad injection. Endotoxin levels were quantified using the Pierce Limulus Amebocyte Lysate Chromogenic Endotoxin Quantitiation kit (ThermoScientific) to be less than 0.5EU/mg for either ovalbumin or ovalbumin conjugated to psDNA.

### Nystatin

Nystatin (Sigma N4014) was resuspended in DMSO to a concentration of 10 mg/mL Mice were injected with 50 μL of 10 mg/mL nystatin per footpad one hour prior to injection with ovalbumin conjugated to Alexa 488 (5 μg) in a mixture with polyI:C and anti-CD40 (2.5 μg each). LNs were harvested and digested as below (preparation of single cell suspensions) and stained with CD45 brilliant violet 510 (Biolegend clone 30F11, 1:300), PDPN APC (Biolegend clone 8.1.1, 1:200), CD31 PercP Cy5.5 (Biolegend clone 390, 1:200) and PD-L1 pacific blue (Biolegend clone 10F.9G2, 1:200).

### Tetramer and intracellular cytokine assays

Draining LNs were processed by glass slide maceration 7 days after injection, washed and suspended in FACS (2% FBS in PBS) buffer containing **Tetramer** (SIINFEKL)-PE (1:400) (NIH tetramer core facilty), **CD8** APC-Cy7 (Biolegend clone 53-6.7 1:400) for 1hr at 37C. Cells were washed and stained for 30 minutes in **CD44** PerCP Cy5.5 (Biolegend clone IM7, 1:400), **B220** BV510 (Biolegend clone RA3-6B2, 1:300). Samples were ran on the FACs Canto II flow cytometer (BD).

### Preparation of single-cell suspensions

Two days or two weeks following vaccination with 1E3 CFU of VV-WR with 10μg of ova-psDNA per foot pad, popliteal LNs were removed from 15 mice and LNs were pulled-apart with 22-gauge needles. Tissue was digested with 0.25mg of Liberase DL (Roche, Indianapolis, IN) per ml of EHAA media with DNAse (Worthington, Lakewood, NJ) at 37 degrees. Every 15 minutes media was removed, cells spun down and new digestion media added to the undigested tissue until no tissue remained, ~1hr. Following digestion cells were filtered through a screen and washed with 5mM EDTA in EHAA. LN cells were then divided into thirds where one third underwent staining with CD11c (N418), CD11b and B220 and a live/dead dye (Tonbo). Live cells were then sorted into 4 tubes on a FACs Aria Cell Sorter (BD): sorted CD11c-APC Cy7 (Biolengend clone N418 1:400)+ cells, sorted CD11b PE-Cγ7 (Biolegend clone M1/70)+ cells, sorted B220 PE (Biolegend clone RA3-6B2)+ cells and Fixable Viability Stain 510 (BD Biosciences Cat # 546406) ungated live cells which were recombined at a 4:4:1:1 ratio, respectively. For the remaining two-thirds of cells, cells were stained with CD45 PE followed by magnetic bead isolation using the Miltenyi bead isolation kit. CD45 negative cells that passed through the column were then washed. Both sorted and selected (CD45+ and CD45-) cells were then washed with PBS in 0.1% BSA as described in the Cell Prep Guide (10x Genomics) and counted using a hemacytometer. Final concentration of cells was approximately 1000 cells/ul and approximately 10-20 μL were assayed.

### Single-cell library preparation using the 10x Genomics platform

Cells were assayed using the 10x Genomics single-cell 3’ expression kit v3 according to the manufacturer’s instructions (CG000183 Rev B) and CITE-seq protocol (cite-seq.com/protocol Cite-seq_190213) with the following changes:

1. *cDNA amplification and cleanup*. During cDNA amplification, 1 μL of 0.2 uM each mixture of additive forward and reverse primers (**Supplementary Table 1**) was included to amplify the antigen tags. The CITE-seq protocol was followed for size selection and clean up of the cDNA and antigen tag products. Antigen tag products were eluted in 60 μL of nuclease-free water.
2. *Amplification of antigen tag sequencing libraries*. A 100 μL PCR reaction was prepared containing 45 μL of purified antigen tag products, 1X Phusion HF Buffer (NEB), 200 uM dNTPs, 25 pmol each Illumina sequencing forward and reverse primers (**Supplementary Table 1**), 2 Units Phusion High Fidelity DNA Polymerase. PCR cycling conditions were 95 °C for 3 min, 6-10x(95 °C for 20 s, 60 °C for 30 s, 72 °C for 20 s), 72 °C for 5 min. Products were purified according to the CITE-seq protocol. *G*ene expression and antigen tag libraries were analyzed on the Agilent D1000 Tapestation and quantified using the Qubit HS dsDNA fluorometric quantitation kit (Thermo Scientific).

All libraries were sequenced on a Illumina NovaSeq 6000 with 2 x 150 base pair read lengths.

### Transcriptome and oligonucleotide detection and analysis

Briefly, FASTQ files from the gene expression and antigen tracking libraries were processed using the feature barcode version of the cellranger count pipeline (v3.1.0). Reads were aligned to the mm10 and Vaccinia virus (NC_006998) reference genomes. Analysis of gene expression and antigen tracking data was performed using the Seurat R package (v3.2). Antigen tracking and gene expression data were combined into the same Seurat object for each sample (CD45-/day2, CD45+/day2, CD45- /day14, CD45+/day14). Cells were filtered based on the number of detected genes (>250 and <5000) and the percent of mitochondrial reads (<15%). Gene expression counts were log normalized (NormalizeData), and relative ova signal was calculated by dividing ova-psDNA counts by the median ova-psDNA counts for all T and B cells present in the sample. Gene expression data were scaled and centered (ScaleData). 2000 variable features (FindVariableFeatures) were used for PCA (RunPCA) and the first 40 principal components were used to find clusters (FindNeighbors, FindClusters) and calculate uniform manifold approximation and projection (UMAP) (RunUMAP). Cell types were annotated using the R package clustifyr (https://rnabi-oco.github.io/clustifyr) ^48^ along with reference bulk RNA-seq data from ImmGen (available for download through the clustifyrdata R package, https://rnabioco.github.io/clustifyrdata). The samples were then divided into separate objects for DCs, LECs, and FRCs. Cell subsets were then annotated using clustifyr with reference bulk RNA-seq data for DCs^49, 50^, FRCs^51^, and LECs^52, 53, 54^.

Identification of ova-low and -high populations was accomplished using a two-component Gaussian mixture model implemented with the R package mixtools (https://cran.r-pro-ject.org/web/packages/mixtools/index.html). All LECs were used when identifying ova-low and ova-high cells (**Fig. 4**). For DCs (**Supplementary Fig. 10**), ova-low and -high populations were identified independently for each DC cell type. For ova-low and ova-high populations, differentially expressed genes were identified using the R package presto (wilcoxauc, https://github.com/immunogenomics/presto). Differentially expressed genes were filtered to include those with an adjusted p-value <0.05, log fold change >0.25, AUC >0.5, and with at least 50% of ova-high cells expressing the gene.

### Raw data and analysis software

Raw and processed data for this study have been deposited at NCBI GEO under accession GSE150719. A reproducible analysis pipeline is available at https://github.com/rnabioco/antigen-tracking.

### Statistical analysis

Statistical analysis was done using either a non-parametric 2 tailed Mann-Whitney t-test or multiple t-tests with a two stage step up up method of Benjamini, Krieger and Yekutieli without assuming consistent standard deviations. Each in vivo analysis was performed with 3-6 mice per group as determined by a power calculation using the assumption (based on prior data) that there will be at least a two fold change with a standard deviation of less than 0.5. To calculate numbers we performed a power calculation with an alpha of 0.5 and a 1-beta of 0.80 to determined at least 3 mice per group should be evaluated.

## ACKNOWLEDGEMENTS

The K^b^ SIINFEKL PE tetramer was provided by the NIH Tetramer Core Facility. We thank the HIMSR flow cytometry core facility for use of the Aria cell sorter and University of Colorado Anschutz Medical Campus Genomics Core Facility (NIH P30 CA046934).

## AUTHOR CONTRIBUTIONS

S.M.W, T.D, E.D.L., M.A.B and B.A.J.T. performed experiments. S.M.W, T.D, E.D.L., M.A.B, R.M.S., R.F. and B.A.J.T. analyzed the data from the experiments. B.A.J.T., S.M.W., T.D., J.R.H. designed the experiments. B.A.J.T, S.M.W., and J.R.H. wrote the paper.

---

**Supplemental Figure 1:**
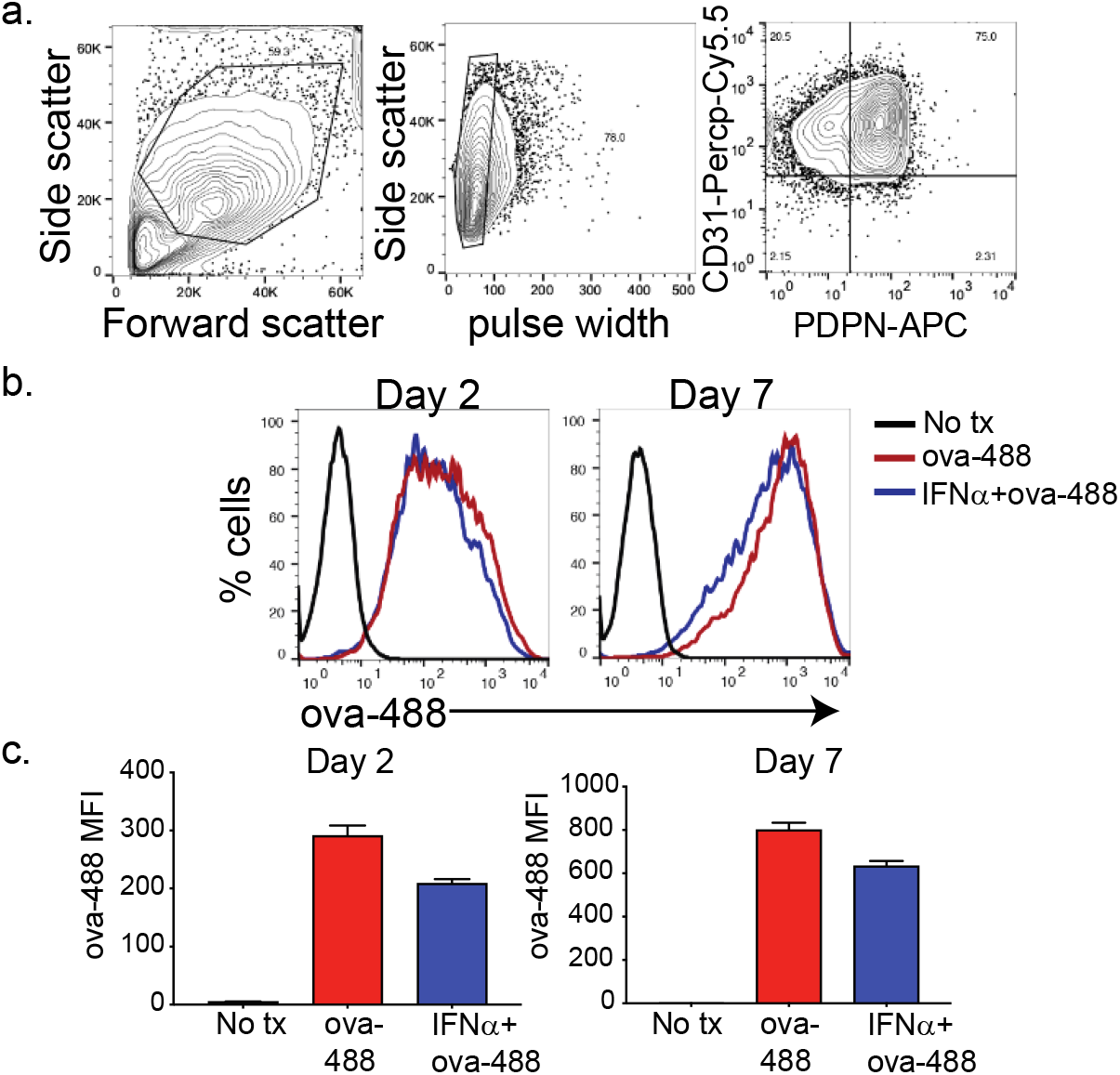
**A**. Gating strategy for murine lymphatic endothelial cells. **B**. Amount of ovalbumin conjugated to alexafluor 488 that was acquired over a two to seven day period in the presence or absence of type 1 IFN. **C**. Quantification of fluorescence intensity two or seven days after treatment as indicated.

**Supplemental Figure 2:**
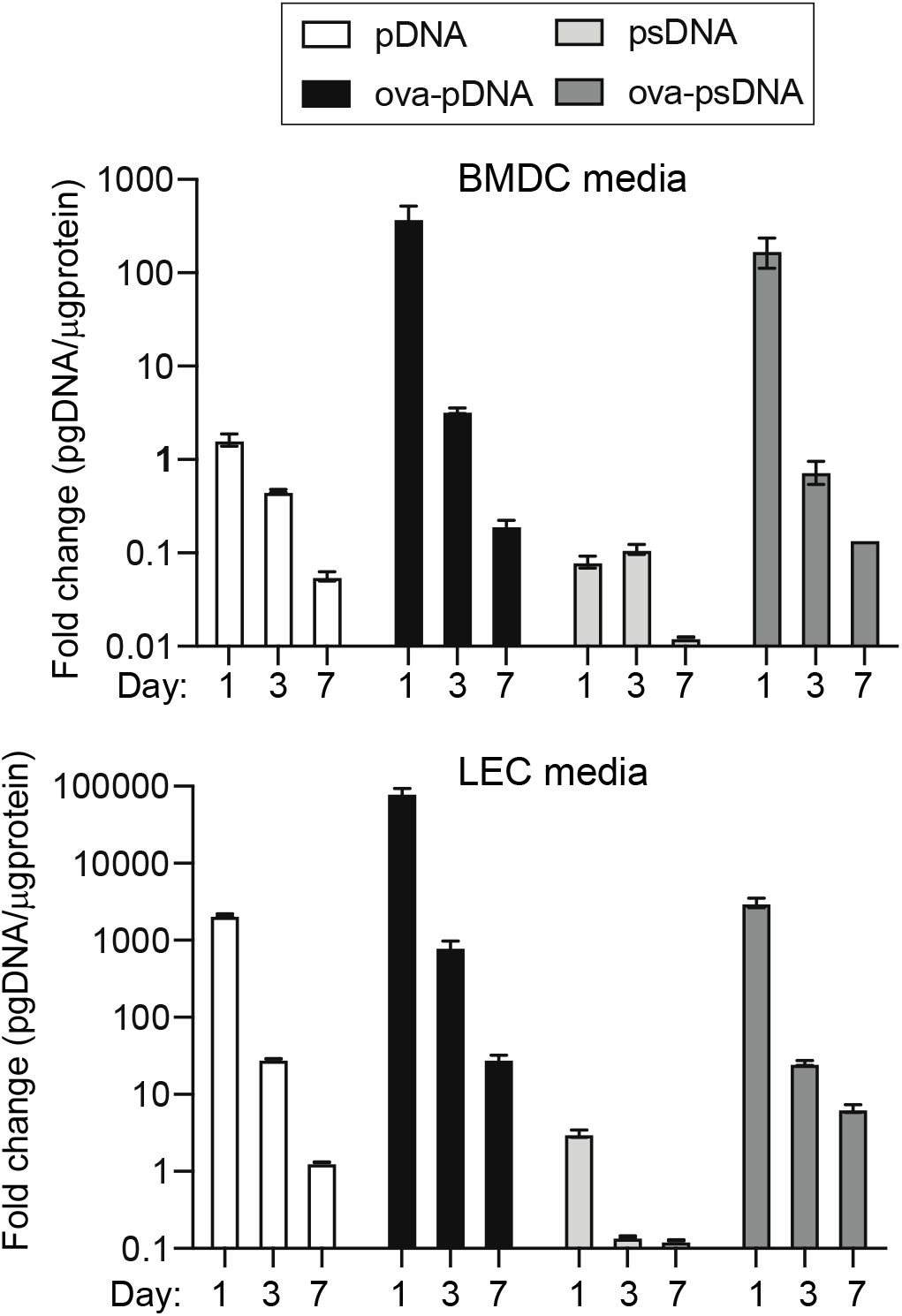
DNA barcode is not retained in cell media over time. **A**. Bone marrow derived dendritic cells (BMDC) were grown for 7 days and cultured in GM-CSF. After 7 days 5μg of either pDNA, psDNA, ova-pDNA or ova-psDNA were added to the culture media and 1,3, or 7 days after addition, media was removed and qPCR was performed on using primers against the DNA added. **C**. As in B except with murine lymph node lymphatic endothelial cells.

**Supplemental Figure 3.**
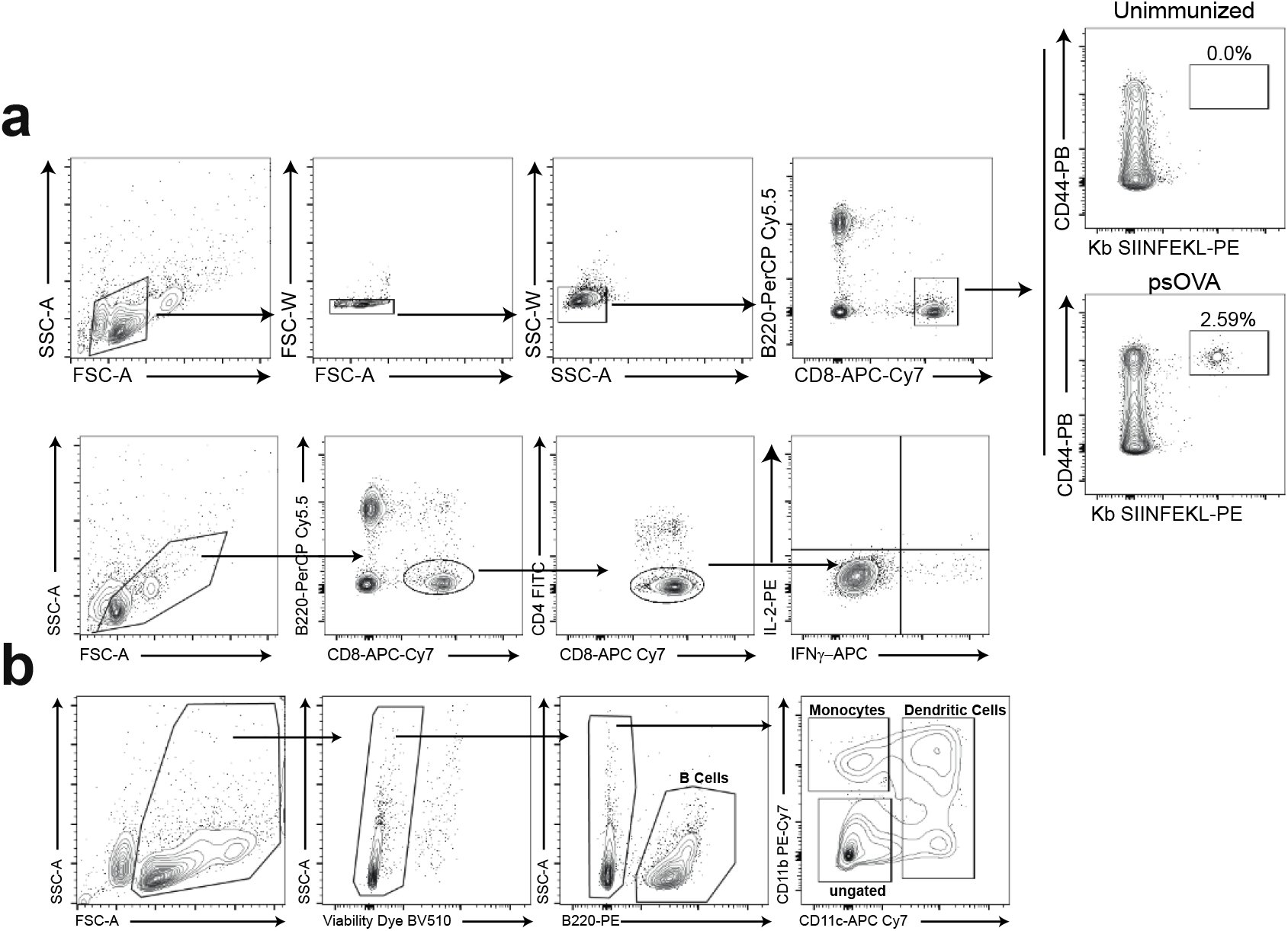
Gating strategies. **A**. Gating strategy for tetramer staining and intracellular cytokine staining of CD8 T cells. **B**. Gating strategy used for cell sorting prior to single cell RNA sequencing.

**Supplemental Figure 4.**
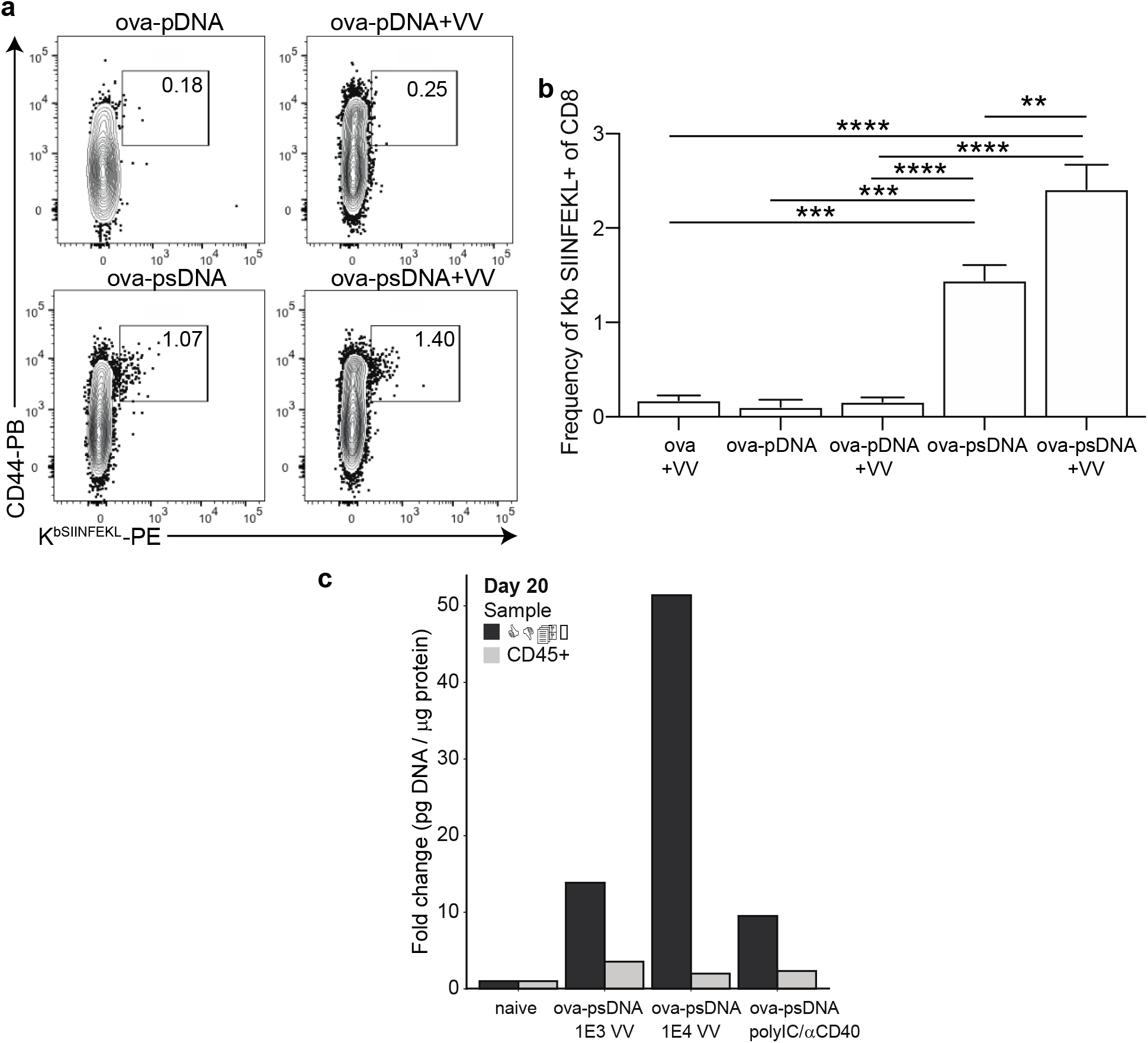
Vaccinia plus ova-DNA conjugate induces ova specific T cell response and archiving. **A**. Mice were vaccinated in the footpad with 10 μg of ovalbumin-pDNA (phosphodiester backbone) or ovalbumin-psDNA (phosphorothioate backbone) with or without vaccinia virus (10_3_ PFU). Each barcode conjugate contained a unique sequence barcode. Seven days after vaccination, mice were euthanized and draining popliteal LN cells were isolated and stained. Shown are B220-, CD3+, CD8+ cells. Box indicates the frequency of tetramer specific and CD44 high cells per lymph node. **B**. Quantification of A. **C**. Mice were vaccinated with 10μg ovalbumin conjugated to phosphorothioate DNA (ova-psDNA) plus 1E3 VV-WR, 1E4 VV-WR or polyI:C/aCD40 (5μg each) in each footpad. LNs were harvested twenty days later and miltenyi bead selection was performed using CD45 to select hematopoietic (CD45+) versus non-hematopoietic (CD45-) cells. The amount of DNA barcode was assessed in each group as a faithful reporter of antigen archiving. Values aer displayed as fold-change relative to the negative control naïve sample.

**Supplemental Figure 5.**
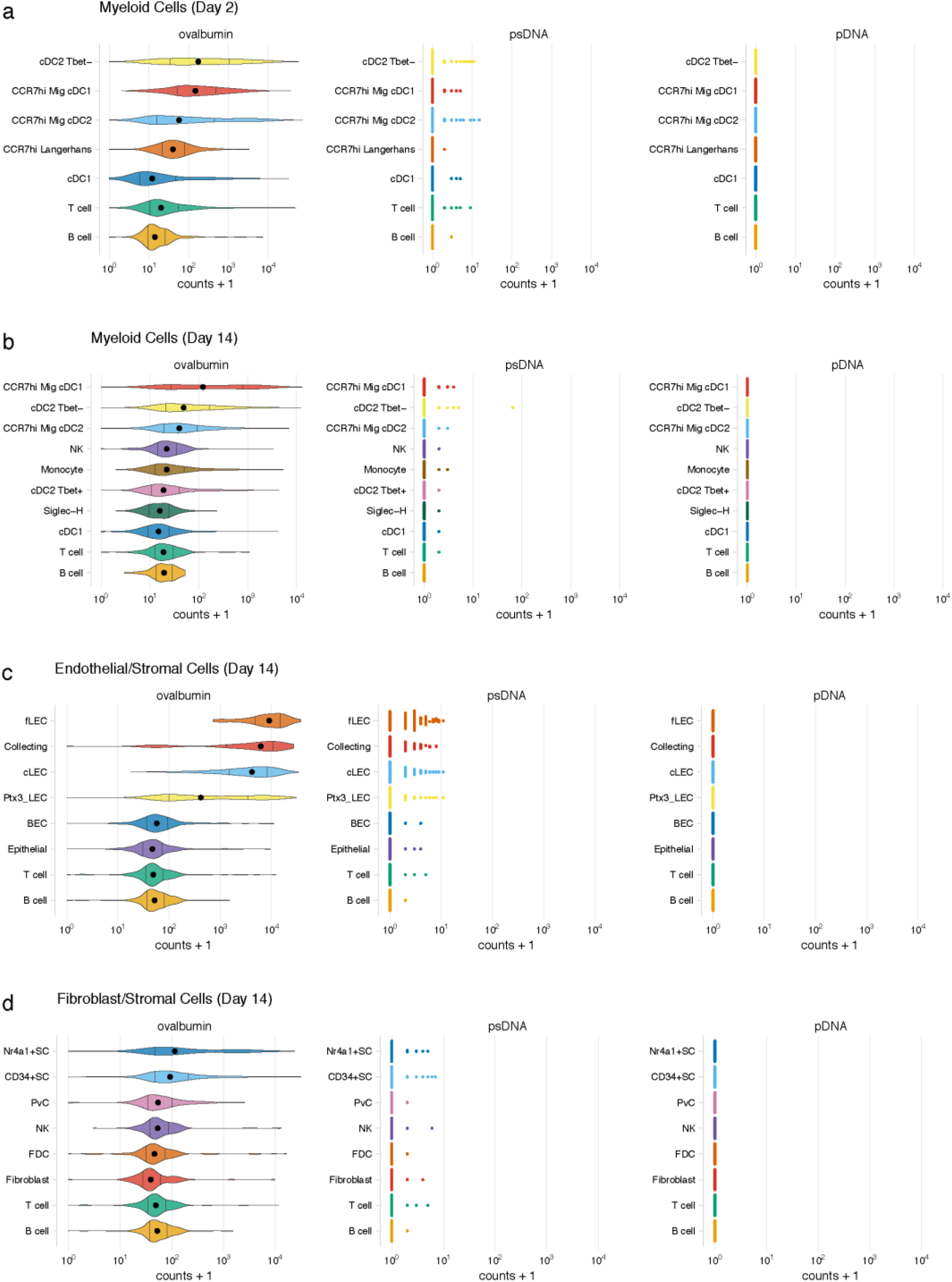
Detection of DNA barcode requires conjugation to ovalbumin. Mice were vaccinated with ova-psDNA, psDNA and pDNA with IE3 CFU of VV-WR. Each DNA injected had a unique barcode sequence for detection during sequencing. Counts of ova-psDNA, psDNA, pDNA for DCs at A. day 2 or B. day 14. Counts for ova-psDNA, psDNA and pDNA for C. LECs and D. FRCs 14 days post vaccination.

**Supplemental Figure 6.**
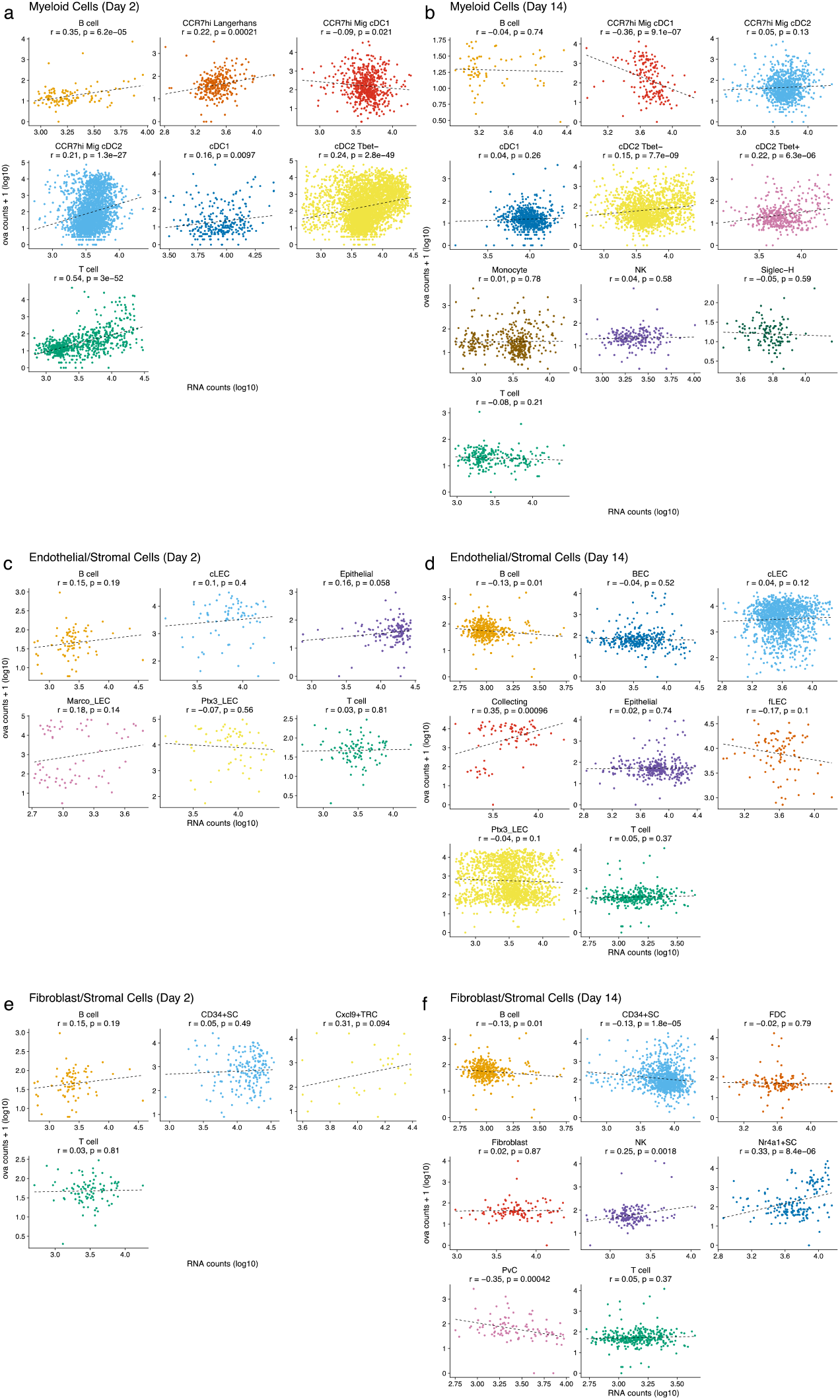
Antigen counts were independent of total mRNA counts. Antigen counts were compared with total mRNA counts for each cell for DCs (A,B), LECs (C,D) and FRCs (E,F). Pearson’s correlation coefficient and associated p-value is shown for each cell type.

**Supplemental Figure 7.**
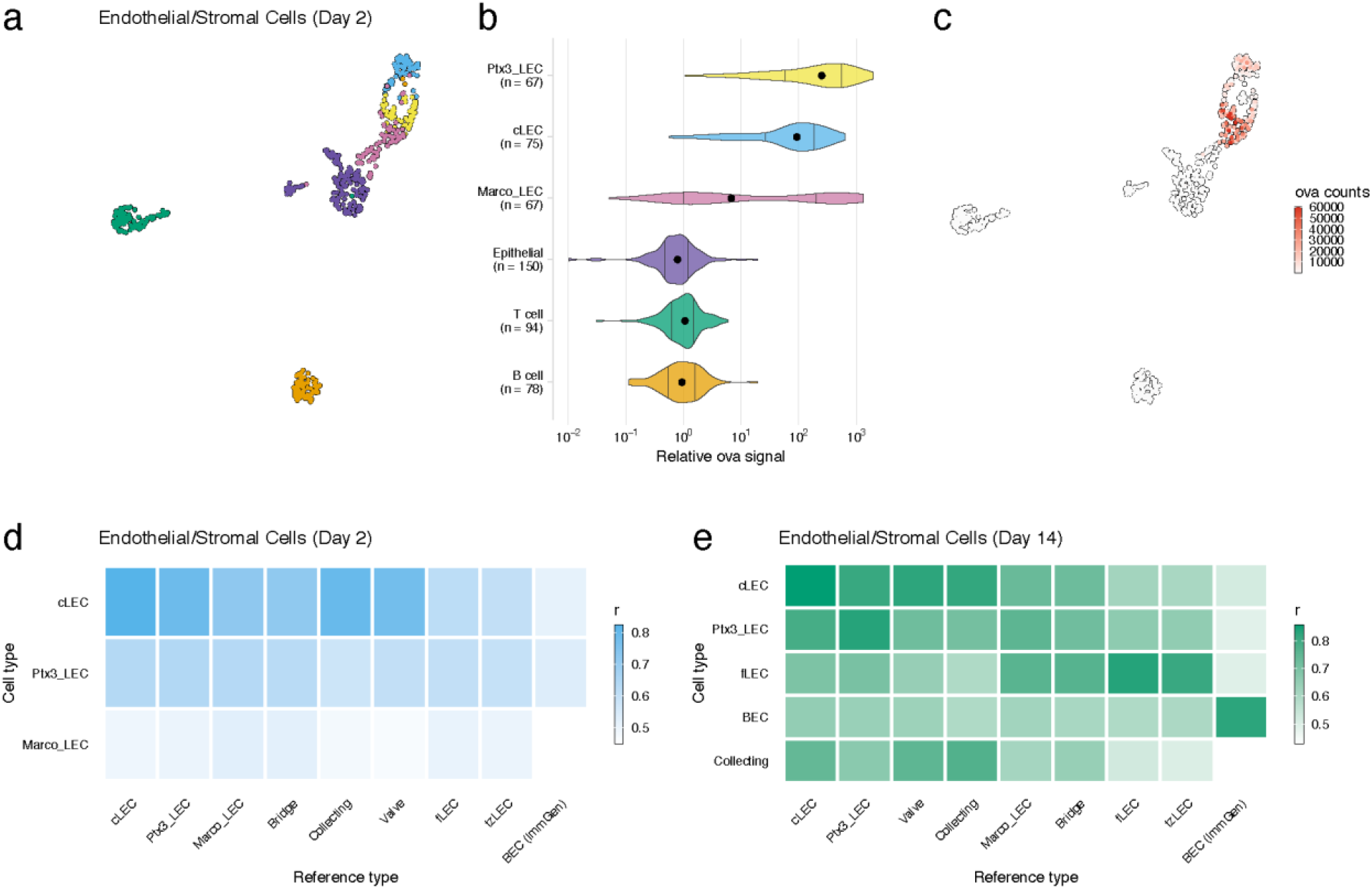
LEC cell types associated with high antigen counts 2 days after vaccination. A. A UMAP projection is shown for LEC cell types, epithelial cells, B cells and T cells identified for the day 2 timepoint. B. Relative ova signal is shown for each cell type. Relative ova signal was calculated by dividing antigen counts for each cell by the median antigen counts for T and B cells. C. Antigen counts are displayed on the UMAP projection shown in A. D,E. Correlation coefficients are shown comparing each identified LEC cell type with the reference cell types from Xiang et al.

**Supplemental Figure 8:**
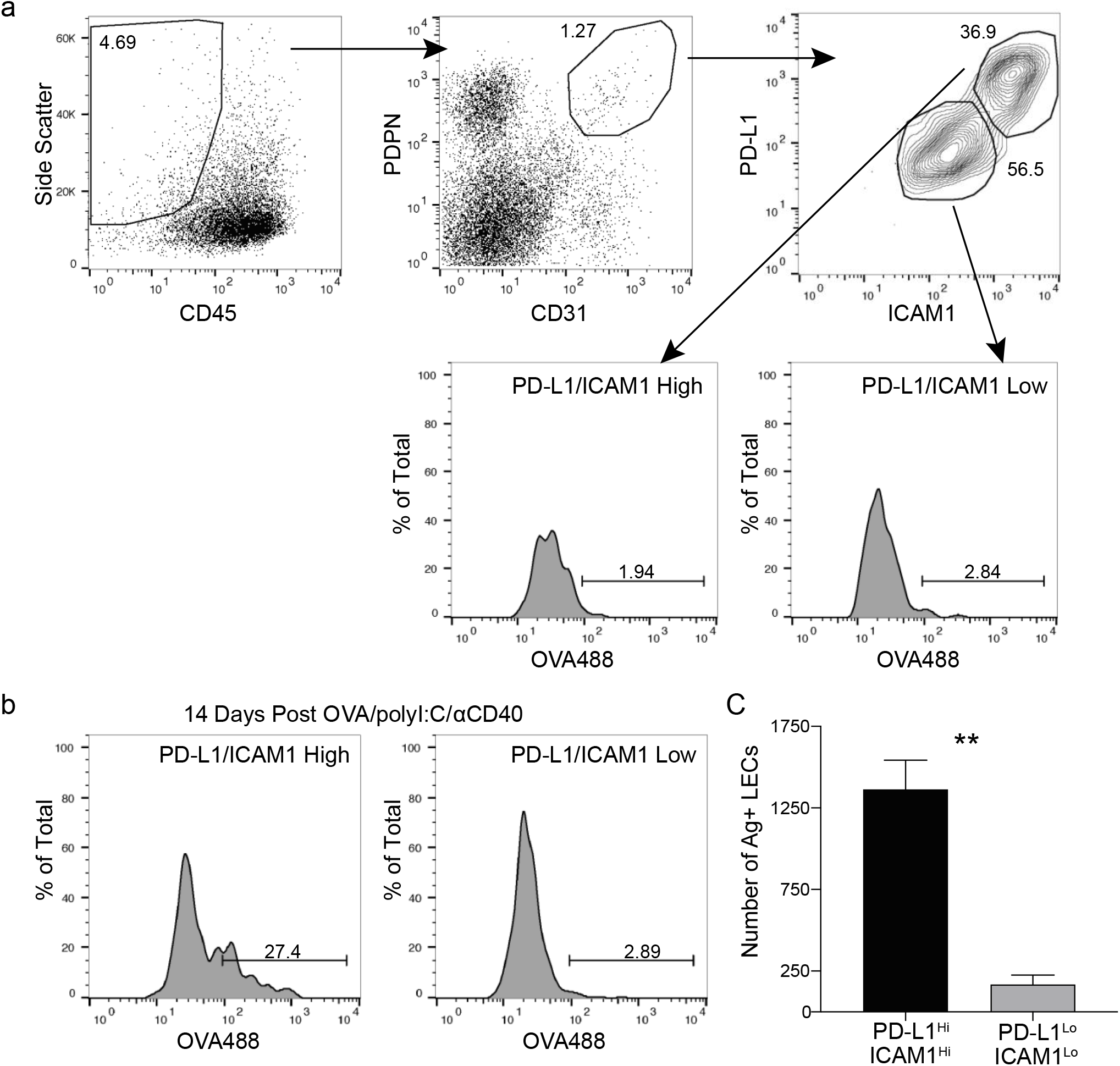
Antigen is held by PD-L1/ICAM1 High LECs. A) Gating strategy for LECs in the dLN with markers for ICAM1 and PD-L1. B) Representative flow plots of antigen held by LECs 14 days following subcutaneous immunization with OVA488 (10μg/site), polyI:C (5μg/site) and αCD40 (5μg/site). LECs were gated first on PD-L1 high/ICAM1 high or PD-L1 low/ICAM1 low and then gated on antigen positive. C) Quantification of B showing the number of antigen positive LECs gated on PD-L1 high/ICAM1 high or PD-L1 low/ICAM1 low. Statistical analysis was done using an unpaired student’s t test, ** p=0.0032

**Supplemental Figure 9.**
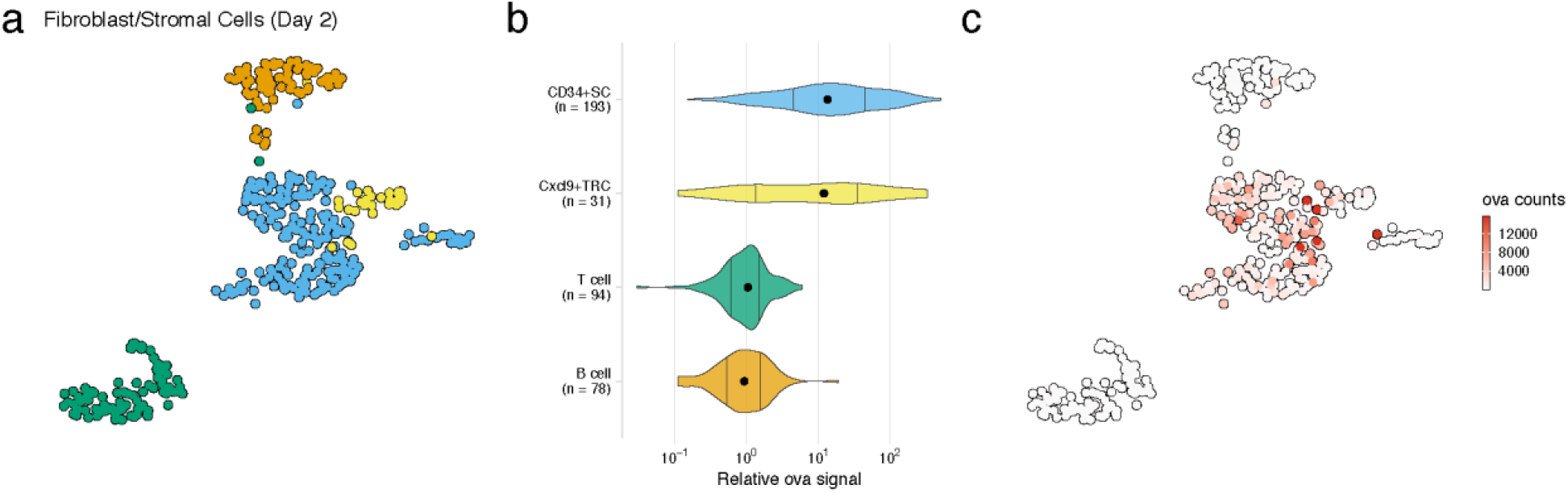
FRC cell types with high antigen counts at day 2 post vaccination. A. UMAP projection is shown for FRC cell types, B cells, and T cells identified for the day 2 timepoint. B. Relative ovalbumin signal is shown for each cell type. Relative ovalbumin signal was calculated by dividing antigen counts for each cell by the median antigen counts for T and B cells. C. Antigen counts are displayed on the UMAP projection shown in A.

